# Structural basis of flagellar filament asymmetry and supercoil templating by *Leptospira* spirochete sheath proteins

**DOI:** 10.1101/2022.03.03.482903

**Authors:** Megan R. Brady, Fabiana San Martin, Garrett E. Debs, Kimberley H. Gibson, Azalia Rodríguez, Rosario Durán, Elsio A. Wunder, Albert I. Ko, Alejandro Buschiazzo, Charles V. Sindelar

## Abstract

Several Leptospira species are bacterial agents of leptospirosis, a neglected tropical disease responsible for ^~^1 million cases and 50,000 deaths each year worldwide. *Leptospira*, like other members of the Spirochaeta phylum, possess specially adapted flagella that remain confined within the periplasm. These appendages drive a unique, corkscrew-like swimming style that enables efficient motility and pathogenesis. However, the composition, function, and molecular architecture of spirochetal flagellar filaments remain poorly understood. We solved single-particle cryo-EM structures of isolated *Leptospira* flagellar filaments, comparing the wild-type form to two mutant forms with different missing components and abrogated motilities. The structures reveal a complex proteinaceous sheath surrounding a conserved core composed of the FlaB flagellin homolog. Sheath proteins were found to fall into two distinct categories, both of which are required for motility. Filament ‘coiling’ proteins, FcpA and FcpB, exert force on the filament when they bind its surface, causing the filament to stretch. In contrast, we identify sheath components FlaAP (newly discovered in this study) and FlaA2 as ‘template’ factors, which have little effect on filament shape by themselves, but partition the coiling proteins to one side of the filament. In this way, the two types of *Leptospira* sheath factors operate collectively on the flagellar filament to bend it from a ‘relaxed’ form associated with cell immobility, to a motility-competent shape that is tightly supercoiled. Our structures also indicate that core-sheath interactions are largely mediated by carbohydrate moieties from flagellin core side chain *O*-glycosylations. The supercoiling mechanism presented here provides a benchmark for studies with other bacteria, for which near-atomic resolution structures of flagellar filament in native supercoiled forms, are still lacking.

## Introduction

Spirochetes constitute an ancient Gram-negative bacterial Phylum that comprises important human and animal pathogens, such as *Borrelia burgdorferi* (the etiologic agent of Lyme disease), *Treponema pallidum* (syphilis), and several species of *Leptospira* (which cause leptospirosis) (Paster 2010). Spirochetes have a spiral-shaped cell body and exhibit unique wavy or ‘corkscrew-like’ swimming motility, allowing them to drill through tissues and highly viscous environments with very high efficiency (Li, Motaleb et al. 2000). This form of locomotion is critical to their pathogenicity, as motility-deficient *Leptospira* are unable to infect their hosts (Lambert, Picardeau et al. 2012, Fontana, Lambert et al. 2016, Wunder, Figueira et al. 2016), making motility mutants attractive vaccine candidates (Wunder, Adhikarla et al. 2021).

Most bacteria generate translational motility through flagellar gyration, powered by rotation of the flagellar motor; torque is then transmitted through a hook to the long flagellar filament that ultimately provokes thrust (Berg and Anderson 1973, Berg 2003). In most bacteria, such as *Salmonella, Campylobacter, Bacillus*, and many others, these flagella are extracellular (Pead 1979, Namba and Vonderviszt 1997). In contrast, Spirochetes contain periplasmic flagella, wrapping around their cell body while completely confined between the inner and outer cytoplasmic membranes (Wolgemuth 2015).

The spirochete *Leptospira* has only one flagellum at both cell poles, with filaments ~3 – 5 μm long that do not overlap towards the center of the cell body (Nauman, Holt et al. 1969, Paster 2010, Picardeau 2017). Clockwise rotation of the flagellar motor distorts the end of the *Leptospira* cell body into a “hook” shape, while counter-clockwise rotation enforces instead a “spiral” shape. Translation will only occur if the two motors are rotating in these opposite directions, in which case the cell will move in the direction of the spiral end, exhibiting a gyrating hook towards the trailing end (Goldstein and Charon 1988, Goldstein and Charon 1990, Wolgemuth, Charon et al. 2006, Wolgemuth 2015). When purified, flagella from wild-type *Leptospira* spontaneously adopt a strong supercoiled conformation, resulting in a flat spiral spring architecture when observed with transmission electron microscopy (Bromley and Charon 1979, Trueba, Bolin et al. 1992, Gibson, Trajtenberg et al. 2020).

Most bacterial filaments are homopolymers that self-assemble from a single protein species, e.g. *Salmonella* flagellar filaments are composed of repeating subunits of the protein flagellin (FliC) (Berg 2003). In contrast, spirochete filaments are composed of several proteins: (i) FlaB, expressed as one or more isoforms in different spirochetes, constitutes the “core” of the appendage, orthologous to the all-helical D0 and D1 domains of FliC (Mitchison, Rood et al. 1991, Lin, Surujballi et al. 1997); and (ii) FlaA isoforms, thought to constitute a proteinaceous sheath that covers the filament core, and not exhibiting detectable homology to flagellar proteins from other bacteria (Brahamsha and Greenberg 1989, Ge and Charon 1997, Li, Corum et al. 2000, Wolgemuth, Charon et al. 2006).

Focusing on *Leptospira*, the flagellar filament contains four FlaB isoforms (FlaB1-4) and two FlaA isoforms (FlaA1, FlaA2) (Malmstrom, Beck et al. 2009, Lambert, Picardeau et al. 2012). *Leptospira* also express at least two additional sheath proteins not found in other spirochetes: Flagellar Coiling Proteins FcpA and FcpB, which contribute to the coiled shape of wild-type filaments (Wunder, Figueira et al. 2016, Wunder, Slamti et al. 2018). As in other bacterial flagella (Wyss 1998, Kurniyati, Kelly et al. 2017, Blum, Filippidou et al. 2019, Kreutzberger, Ewing et al. 2020, Montemayor, Ploscariu et al. 2021), it is presumed that the *Leptospira* filament core is glycosylated (Holzapfel, Bonhomme et al. 2020).

In addition to the pathogenic species of *Leptospira* (e.g., *L. interrogans*), there are also saprophytic species (e.g., *L. biflexa*). The pathogenic and saprophytic strains share ~61% of genes, including all known flagellar genes (Picardeau, Bulach et al. 2008, Evangelista and Coburn 2010, Fouts, Matthias et al. 2016, Wunder, Figueira et al. 2016, Wunder, Slamti et al. 2018). As the saprophytic strains are easier to manipulate genetically and grow faster in culture (Picardeau, Bulach et al. 2008), *L. biflexa* has been used for some previous flagellar studies (Picardeau, Bulach et al. 2008, Sasaki, Kawamoto et al. 2018, Wunder, Slamti et al. 2018, Gibson, Trajtenberg et al. 2020).

Recently, we reported a 3D reconstruction of the native flagellar filament from *Leptospira biflexa*, obtained through cryo-subtomogram averaging. This structure revealed a striking asymmetric distribution of flagellar sheath proteins, with FcpA and FcpB localized to the outer curvature face of the coiled filament (Gibson, Trajtenberg et al. 2020). However, structural and functional roles of these and other sheath proteins are not well understood. This is due in part to a lack of atomic-level descriptions of the flagellar filament for *Leptospira* or any other spirochete.

Here we present single particle cryo-EM structures of three variants of *Leptospira* flagellar filaments: the wild-type filament and two mutant forms with specific sheath components deleted, purified respectively from *fcpA*^-^ and *flaA2*^-^ knock-out strains. The structures provide near-atomic resolution descriptions of three previously described sheath proteins (FlaA2, FcpA, and FcpB) *in situ* on the filament. Moreover, we discovered evidence that several additional, previously undescribed proteins also reside in the sheath. Among these was a FlaA2-associated protein (FlaAP), whose structure was solved *in situ*.

Examination of the mutant filament structures revealed that sheath proteins can be divided into two distinct categories. In the *fcpA*^-^ mutant that lacks both FcpA and FcpB (Wunder, Figueira et al. 2016), ‘asymmetric binders’ FlaA2 and FlaAP colocalize in a single row along one side of the filament, leaving the other side bare. In contrast, ‘coiling proteins’ FcpA and FcpB assemble in a lattice on the filament surface and longitudinally stretch the filament.

In the wild-type filament, these sheath proteins coexist as an asymmetric assembly where the FcpA/FcpB lattice is disrupted by FlaA2 and FlaAP. We propose that the distinctive, tightly supercoiled shape of *Leptospira* filaments arises due to the resultant, asymmetric stretching forces exerted by FcpA/FcpB on the filament. This mechanism explains why neither FlaA2 nor the coiling proteins are alone sufficient for motility and pathogenesis. We also describe a previously unsuspected role for glycosylated side chains from the core FlaB proteins, which were found to mediate interactions with the sheath.

## Results

### Sheath proteins FlaA2 and FlaAP form a row localized to the filament inner curvature

The knock-out deletion of the *fcpA* gene results in a viable, yet non-motile *Leptospira* strain that forms flagellar filaments with only one known sheath factor: FlaA (Wunder, Figueira et al. 2016). The lack of FcpA precludes normal FcpB recruitment and hence the *fcpA*^-^ mutant filaments do not form the tight coils seen in purified wild-type flagella (Wunder, Figueira et al. 2016, Sasaki, Kawamoto et al. 2018). This strain may shed light onto the potential role of additional sheath proteins, such as FlaA, in determining filament shape and function. Single-particle analysis cryo-EM was performed on thin flagellar filaments from the *L. biflexa fcpA*^-^ strain yielding a 3D reconstruction at 3.8 Å resolution (Fig. 1 and Table S1). While the filaments were heterogeneous in structure and composition, single particle refinement and classification (Fig. S1A) yielded the near-atomic resolution structure of a single dominant population (Fig. 1B; ~53% of the total number of filament segments). This structure exhibits a pronounced curvature (Fig. 1C). Moreover, two rows of sheath proteins decorate one side of the filament (the inner curvature; Fig. 1C,E), while the rest of the filament is bare. This sheath structure contrasts with wild-type filaments, which are thicker, much more tightly coiled and are enclosed by a much more extensive sheath layer (Gibson, Trajtenberg et al. 2020).

**Figure 1.**
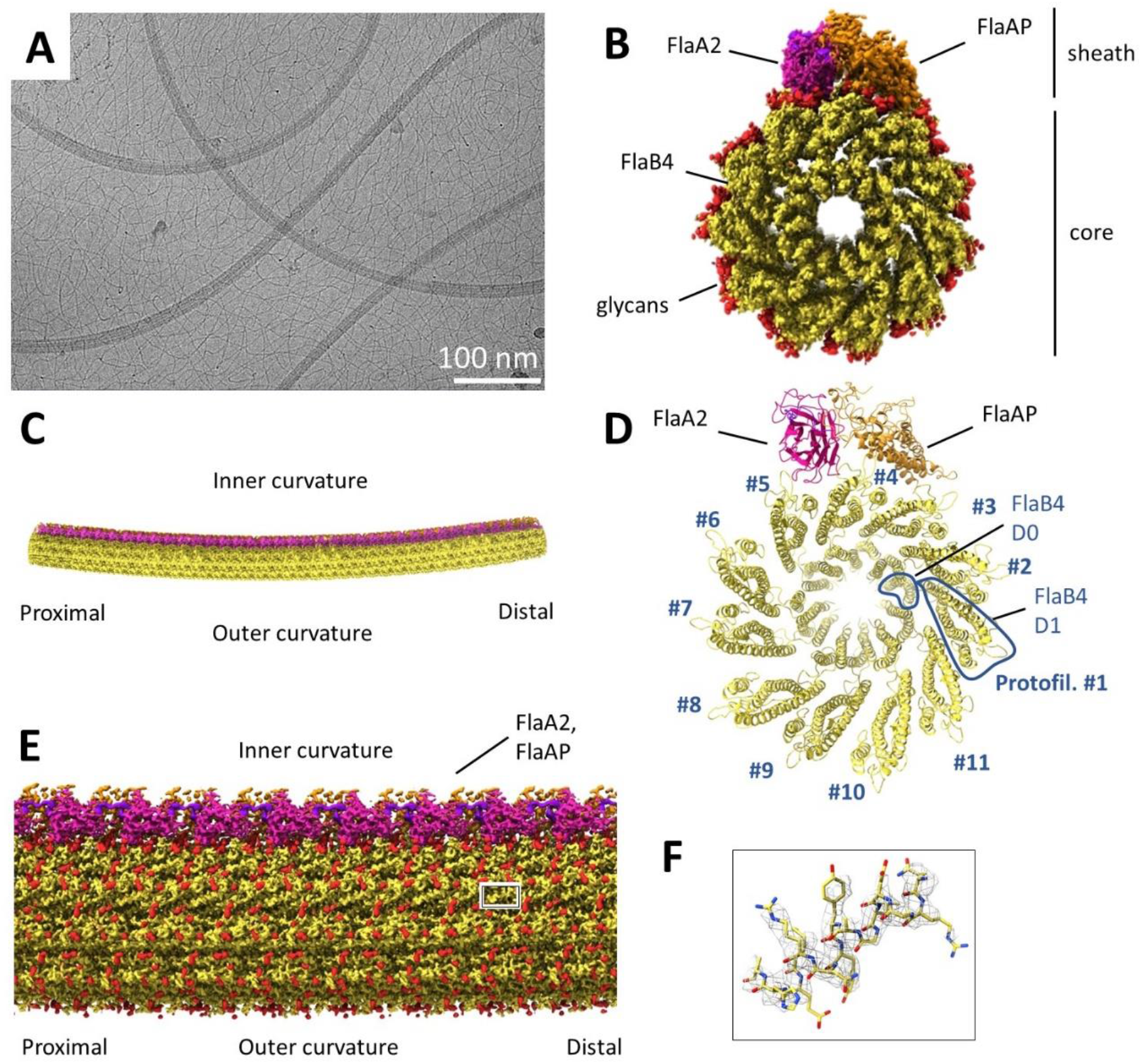
FlaA2 co-localizes to the filament inner curvature in a single row together with FlaAP, a previously uncharacterized protein. In this and the following figures, ‘proximal’ and ‘distal’ labels indicate the direction of the polar filament with respect to the flagellar motor. **A,** Representative micrograph, showing the heterogeneity and slight curvature of the *fcpA*^-^ filaments. Scale bar is 100 nm. **B,** A 3D isosurface rendering of a flagellar filament decorated with FlaA2 and FlaAP. Density for the FlaB core is colored yellow, density identified as bound FlaA2 molecules is colored pink, and density corresponding to FlaAP is colored orange. Putative glycosylation site densities are colored red, and the C-terminal ‘tentacle’ of FlaA2 is colored purple. **C,** A lower magnification view of an extended filament generated from the reconstruction in A, revealing supercoiling. Estimated supercoil pitch and diameter values are 2.01 μm and 0.43 μm respectively. **D,** Models of the FlaB4 core and FlaA2 and FlaAP sheath proteins, using the same coloring as in **B**. The initial models were generated with AlphaFold2, and were fit into the density using Isolde. **E,** Side view of the reconstructed *fcpA*^-^ filament, as in **B**. **F,** A zoomed-in view of the box in **E**, showing electron density corresponding to an alpha-helical segment of FlaB4.

Features in the *fcpA*^-^ density maps were sufficiently well resolved (3.4 – 4.4 Å; Fig. S1C) to build atomic models for the core region composed of the FlaB flagellin homolog, as well as for two distinct sheath proteins: FlaA2, a conserved spirochete sheath factor that was known to be present in this sample (Fig. 1B,D; Fig. 2A,B), and a second, unexpected protein fold that did not match any of the known sheath components of *Leptospira* (Fig. 1B,D; Fig. 2C,D).

**Figure 2.**
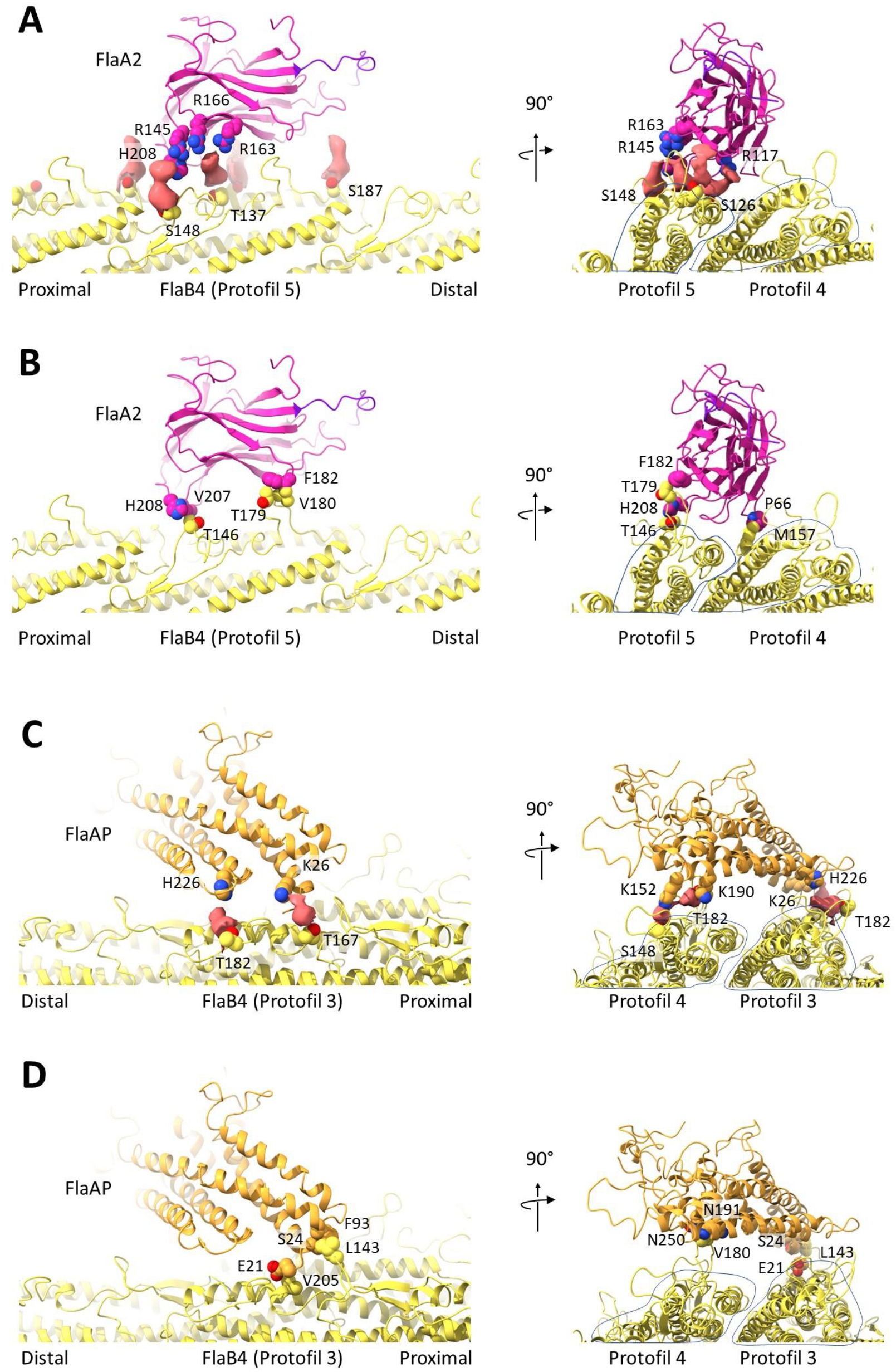
Core interactions of FlaA2 and FlaAP are primarily mediated by glycosylated FlaB4 side chains. **A,** Fit atomic model of FlaA2 highlighting probable sugar-binding sites. Proposed glycan moieties are not modeled, their segmented cryoEM densities are shown as solid pink surfaces. Left: The presumed glycan of Ser_148_ (FlaB4) interacts with Arg_145_ and His_208_ (FlaA2), and the presumed glycan on Thr_137_ (FlaB4) interacts with Arg_163_ and Arg_166_ (FlaA2). Right: End-on view of the filament shows that the glycans are all located along one protofilament (#5). **B,** Protein-protein interactions between FlaA2 and the FlaB4 core. Left: His_208_ and Val_207_ (FlaA2) interact with Ser_146_ (FlaB4), and Phe_182_ (FlaA2) interacts with Thr_179_ and Val_180_ (FlaB4). End-on view, showing that these protein contacts bridge between two adjacent FlaB4 protofilaments (#4 and #5). **C,** Interactions between the FlaAP sheath protein and the glycans associated with the FlaB4 core. Left: The interaction between the presumed glycan on Thr_182_ (FlaB4) and His_226_ (FlaAP), and well as interactions between the presumed glycan of Thr_167_ (FlaB4) and Lys_126_ (FlaAP). Right: End-on view of the filament shows that these glycan interactions bridge between FlaB protofilaments, with two glycan interactions along both protofilaments #3 and #4. **D,** Protein-protein contacts between FlaAP and the FlaB4 core. Left: Ala_172_ (FlaB4) interacts with Glu_21_ (FlaAP), and Leu_143_ (FlaB4) interacts with Phe_93_ and Ser_24_ (FlaAP). Right: End-on view shows that the protein-protein contacts also help to bridge FlaAP between protofilaments #3 and #4.

While neither of these sheath proteins have previously been structurally characterized experimentally, FlaA2 could be identified based on excellent correspondence between its density features and a structure generated by the AlphaFold2 prediction software (Fig. S1D) (Jumper, Evans et al. 2021). Lower resolution limited the ability to trace and sequence the second sheath component directly from the cryo-EM maps. In order to pinpoint the identity of this second component, a mass spectrometry-based proteomics approach was followed, analyzing purified *fcpA*- flagellar filaments (Table S2; see Methods). Approximately 100 uncharacterized proteins were thus identified in triplicate replicas from independent purifications, apart from the known filament proteins. The 3D structures of the five most abundant uncharacterized proteins were predicted with AlphaFold2, confirming that the most abundant species explained the electron density._We have named this second sheath component FlaAP (for FlaA-associated Protein), and found it to correspond to the hypothetical protein sequence encoded by gene LEPBI_I0551 (Picardeau, Bulach et al. 2008), which is present across all *Leptospira* species.

FlaA2 forms a jellyroll fold, with the concave side of the beta sandwich forming a pocket that faces the flagellar core surface and forms a key part of the core interface. Density for FlaA2 was sufficiently well resolved to confirm sequence-specific side chain features, notably including several positively charged side chains lining the core interface (Fig. S2A; see section on glycan binding below). FlaAP folds into a loosely packed alpha helical bundle, with several long, protruding partially ordered loops. While side chain features were not well resolved in this region, density was observed for all predicted secondary structure features from the AlphaFold2 model, except for one disordered loop.

Adjacent FlaA2 molecules form a single row following a single FlaB protofilament (protofilament #5; Fig. 1D), directly interacting with FlaB protofilaments #4 and #5. FlaAP packs laterally against FlaA2 to form its own row, following the subsequent core protofilament (#4), next to the FlaA2 row. Together, these two proteins account for all visible sheath density in the *fcpA*^-^ structure; unique features of their interactions with each other and with the FlaB core are suggestive of functional roles in recognizing flagellar asymmetry and supercoiling as further described below.

### Supercoiled structure of the FlaB core reveals a ‘seam’

The FlaA2 and FlaAP sheath components lie on the surface of a ~120 Å diameter hollow tube (the ‘core’; Fig. 1B) that broadly conforms to the flagellar filament architecture previously described in *Leptospira* and other bacteria (Yonekura, Maki-Yonekura et al. 2003, Maki-Yonekura, Yonekura et al. 2010, Gibson, Trajtenberg et al. 2020). In *Leptospira* and other spirochetes, the core is composed of one or more isoforms of FlaB, a flagellin homolog (Norris, Charon et al. 1988, Li, Motaleb et al. 2000). Our map resolves the core assembly to 3.4 Å resolution or better (Fig. S1C), visualizing all residues of FlaB and allowing a complete atomic model to be built. As observed in other flagellar structures (Yonekura, Maki-Yonekura et al. 2003, Maki-Yonekura, Yonekura et al. 2010, Wang, Burrage et al. 2017, Blum, Filippidou et al. 2019, Kreutzberger, Ewing et al. 2020, Montemayor, Ploscariu et al. 2021), FlaB in our structure has two distinct subdomains (D0 and D1; Fig. 1D), each composed mainly of an extended bundle of alpha helices. Also as in other structures, D0 and D1 are connected by a linker composed of short loop regions. Individual FlaB molecules assemble into linear arrays (protofilaments), 11 of which pack laterally to form the full core assembly (Fig. 1D).

In contrast to previously reported flagellar filament structures (Yonekura, Maki-Yonekura et al. 2003, Maki-Yonekura, Yonekura et al. 2010, Wang, Burrage et al. 2017, Kreutzberger, Ewing et al. 2020, Montemayor, Ploscariu et al. 2021), which correspond to idealized straight helical assemblies, our *fcpA*^-^ structure is curved and exhibits a pronounced supercoil. The shape parameters measured for our *fcpA*^-^ structure (left-handed supercoil with pitch and diameter of 2.0 μm and 0.4 μm respectively) are distinctly different than those measured for the wild-type filament (pitch and diameter of 0.22 μm and 0.45 μm respectively) (Gibson, Trajtenberg et al. 2020). This reflects a loss in the mutant of the characteristic tight and flattened supercoils of wild-type filaments, distortions that are linked to abrogated cell motility and pathogenicity (Wunder, Figueira et al. 2016). Instead, the supercoiled shape of *fcpA*^-^ filaments approaches the ‘normal’ flagellar form observed in exo-flagellated bacteria such as *Salmonella enterica* and *Escherichia coli* (Fig. S3A) (Leifson 1960, Fujii, Shibata et al. 2008, Wang, Jiang et al. 2012). Detailed structural descriptions of native supercoiled flagellar filaments have previously been unattainable for these or any other bacteria. In addition to informing *Leptospira* motility, the *fcpA*^-^ structure presented here thus provides an important reference for assessing general models of natural bending and supercoiling in bacterial flagella.

Due to supercoiling, strict 11-fold helical symmetry in the core of our *fcpA*^-^ structure is broken, and the inner curvature of the supercoil undergoes specific structural changes, broadly consistent with a ‘polymorphic switching’ model consistent with observations reported on *Salmonella* flagella (Maki-Yonekura, Yonekura et al. 2010, Calladine, Luisi et al. 2013). A longitudinal sliding of ~2 Å was observed between D1 domains of protofilaments #4 and #5 on the inner curvature side of the filament (Fig. S3B). This sliding is associated with a conformational change in protofilament #4 characterized by minor deformations in the D0-D1 linker that bring the D1 proximal end ~2 Å closer to the distal end of D0 (Fig. S3B). The same movement also brings the D1 proximal end closer into contact with D1 of the longitudinally adjacent neighbor (*i*-11) in the same protofilament (Fig. S4).

Each of the above behaviors recapitulates predicted conformational changes for a transition between two main flagellin conformations (‘R’ type and ‘L’ type) that underly the polymorphic switching model. We therefore identify the conformation of FlaB in protofilament #4 as ‘R’ type, while the remaining 10 protofilaments are considered ‘L’ type. We note, however, that flagellar supercoiling behavior in this organism is not yet demonstrated to conform to the polymorphic switching model. Indeed, the polymorphic switch model itself remains incompletely proven (see Discussion).

In our *fcpA*^-^ supercoiled structure, consistent with the polymorphic switching model, the largest structural rearrangements are concentrated along a single protofilament (Fig. S5). This results in a ‘seam’ that breaks the underlying 11-fold helical subunit symmetry of a straight filament during the transition to a supercoil.

### FlaA2 and FlaAP localize to the seam

The FlaA2 sheath protein straddles protofilaments #4 and #5, where the seam is located, making small but significant protein-protein contacts with the D1 domains of each underlying FlaB (Fig. 2B). Because protofilaments #4 and #5 are axially offset from each other compared with the rest of the filament (due to the ‘sliding’), they present a unique binding interface for FlaA2. While the magnitude of the sliding displacement at the seam is relatively small, ~1.7 Å (Fig. S5), the FlaA2-FlaB interaction surfaces are minimal and point-like (as only four FlaA2 side chains contribute; Fig. 2B). FlaA2 interactions at non-seam locations are therefore likely to be disrupted, providing an explanation for why FlaA2 localizes to the seam in our structure.

FlaAP straddles protofilaments #3 and #4 and has a similar footprint on these protofilaments as FlaA2 has on protofilaments #4 and #5 (Fig. 2D). However, unlike FLaA2, FlaAP does not straddle the seam, and its core interface does not overlap with that of FlaA2. Thus, the interface geometry of FlaAP with FlaB is altered compared to that of FlaA2.

### FlaA2 and FlaAP form a lattice interconnected by flexible loops

Neighboring FlaA2 subunits along a protofilament are connected by cable-like features. These cables, or tentacles, correspond to the FlaA2 C-terminus (residues 226-241), which projects from the edge of the jellyroll beta-sandwich pointing towards the distal end of the filament and extends unsupported over ~15Å to bridge the gap between adjacent FlaA2 molecules (Fig. 3C). Residues 231-241 occupy density on the neighboring FlaA2 subunit, with the terminal residue (Trp_241_) located in a hydrophobic pocket within that adjacent monomer. An additional longitudinal contact is formed by a partially ordered loop (residues 178-196) which presents the side chain of Arg_188_ to that of Tyr_144_ in the next distal FlaA2 subunit to form a probable cation-pi interaction (Fig. 3A).

**Figure 3.**
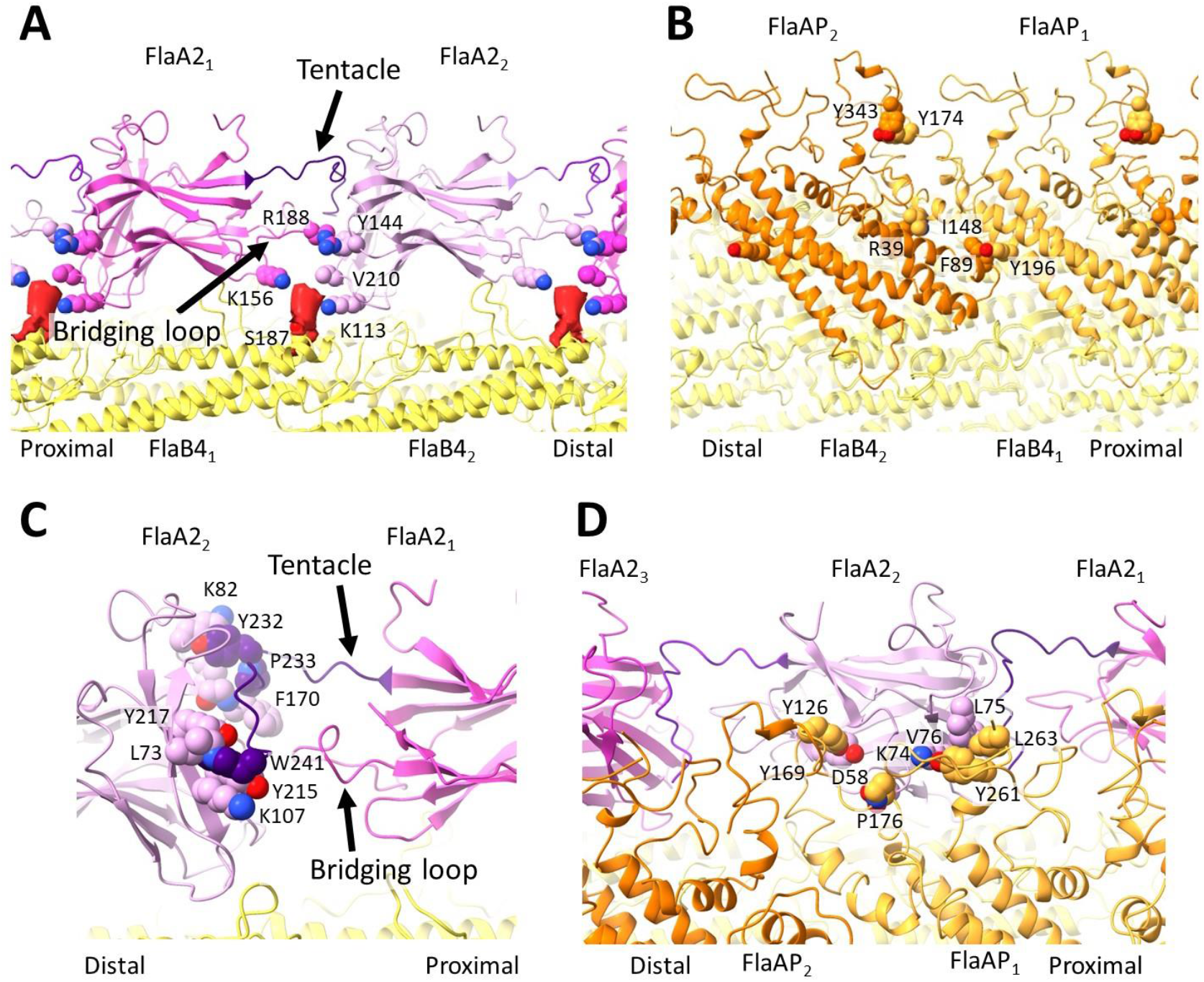
Core interactions of FlaA2 and FlaAP are primarily mediated by glycosylated FlaB4 side chains. **A,** Fit atomic model of FlaA2 reveals probable sugar binding sites. Proposed glycan moieties are not modeled, their segmented cryoEM densities are shown as solid red surfaces. Left: The glycan of Ser_148_ (FlaB4) interacts with Arg_145_ and His_208_ (FlaA2), and the glycan on Thr_137_ (FlaB4) interacts with Arg_163_ and Arg_166_ (FlaA2). Right: End-on view of the filament shows that the glycans are all located along one protofilament (protofilament #5). **B,** Protein-protein interactions between FlaA2 and the FlaB4 core. Left: His_208_ and Val_207_ (FlaA2) interact with Ser_146_ (FlaB4), and Phe_182_ (FlaA2) interacts with Thr_179_ and Val_180_ (FlaB4). End-on view, showing that these protein contacts bridge between two adjacent FlaB4 protofilaments (protofilament #4 and protofilament #5). **C,** Interactions between the FlaAP sheath protein and the glycans associated with the FlaB4 core. Left: The interaction between the presumed glycan on Thr_182_ (FlaB4) and His_226_ (FlaAP), as well as interactions between the presumed glycan of Thr_167_ (FlaB4) and Lys_126_ (FlaAP). Right: End-on view of the filament shows that these glycan interactions bridge between FlaB protofilaments, with two glycan interactions along both protofilaments #3 and #4. **D,** Protein-protein contacts between FlaAP and the FlaB4 core. Left: Ala_172_ (FlaB4) interacts with Glu_21_ (FlaAP), and Leu_143_ (FlaB4) interacts with Phe_93_ and Ser_24_ (FlaAP). Right: End-on view shows that the protein-protein contacts also help to bridge FlaAP between protofilaments #3 and #4.

Longitudinal contacts are also observed between adjacent FlaAP monomers, whose alpha helical bundles directly abut each other (Fig. 3B). Contacts involve only a few side chains, but these tend to be bulky and hydrophobic. An additional longitudinal contact is mediated by a pair of long and flexible loops that pair Tyr_343_ with Tyr_174_ of the next proximal neighbor.

FlaA2 and FlaAP also directly contact each other, via three long and meandering loops of FlaAP. Lower resolution of these loop regions in our map is consistent with greater mobility, but at least two specific contacts are identified in the map, involving clusters of mostly hydrophobic contacts (Fig. 3D). These contacts complete a lattice of nearest neighbor interactions that hold the FlaA2-FlaAP sheath assembly together. As we have described, this lattice is characterized by sparse hydrophobic interactions involving extended and flexible loops. This structural characteristic suggests that the FlaA2-FlaAP assembly may be able to accommodate a variety of filament curvatures.

### FlaA2 and FlaAP interact with glycosylated surface residues of FlaB4

Glycosylated residues on the FlaB core surface were observed to contribute extensively to the core-sheath interfaces for both FlaA2 and FlaAP (Fig. 2A,C). Our map directly visualizes excess densities corresponding to sites of glycosylation (Kurniyati, Kelly et al. 2017, Holzapfel, Bonhomme et al. 2020). These large, elongated densities originate from serine and threonine FlaB side chains, and many of them directly contact the sheath proteins. The latter observation contrasts with previously reported flagellar structures (Echazarreta, Kepple et al. 2018, Kreutzberger, Ewing et al. 2020), where a structural role for glycans has not been observed. While the FlaB glycans in *Leptospira* have not yet been chemically characterized, we infer that they are composed of an extended polysaccharide chain based on (1) their sizes and shapes in our map; and (2) the propensity of these glycans to simultaneously interact with multiple positively charged side chains from their protein partners.

Glycan interactions at the sheath-core interface are dominated by bridging interactions involving positively charged side chains (Lys, His, Arg) from FlaA2 and FlaAP (Fig. 2A,C). Our map identified four such contacts for FlaA2, including a prominent one involving glycosylated Ser_126_. This glycan projects into a shallow pocket formed by the concave beta-sheet face of FlaA2, where it contacts Arg_117_ and Lys_151_ (Fig. 2A). The three remaining glycans surround the periphery of FlaA2 and clasp its exterior, each of them engaging a pair of FlaA2 side chains (Fig. 2A). One of these glycans, at site Ser_187_, also contacts residues on the proximally adjacent FlaA2 subunit, thus bridging the two subunits (Fig. 3A).

Positively charged residues of FlaAP (Lys, His) interact with the glycans of FlaB4 in a manner like that seen with FlaA2. However, while the FlaA2 interactions generally involve two FlaA2 side chains per glycan, for FlaAP we identified only one side chain per glycan interaction (Fig. 2C). Another difference with FlaA2 is that FlaAP uses glycan interactions to straddle two protofilaments in addition to protein-protein interactions (Fig. 2C,D).

Patterns of core glycosylation in our map provide a unique fingerprint, indicating that, of the four FlaB isoforms present in *Leptospira*, FlaB4 predominates. Seven surface-exposed serine and threonine residues of each subunit exhibit signs of glycosylation. Two of these glycosylation sites (Thr_137_ and Thr_182_) are substituted in the other FlaB isoforms by residues which are not amenable to glycosylation (Gln, Glu, Asp at 137 and an Asn, Ile, Glu at 182). Features consistent with the FlaB4 glycosylation fingerprint are conserved across all 11 protofilaments in our asymmetric *fcpA*^-^ reconstruction. These features suggest that the core is predominantly composed of FlaB4 in this *L. biflexa* mutant. Mass spectrometry of the purified *fcpA*^-^ flagella detected a mixed population of FlaB isoforms (Table S2), which would be averaged together in our 3D map. FlaB4 and FlaB1 were identified as the largest isoform populations in this mutant, with a statistically insignificant difference between them (Table S2).

Both sheath protein structures show signs of being able to preferentially bind FlaB4 over the other FlaB isoforms. For FlaA2, of the four glycan contacts, three involve Ser/Thr sites conserved among all four FlaB isoforms, but the remaining site (Thr_137_) is not conserved and is amenable to glycosylation only in FlaB4. This feature suggests that FlaA2 may preferentially recognize FlaB4, an idea that is further supported by the isoform-specific nature of the three FlaA2-FlaB4 protein-protein contacts observed in our structure. For the contact between Val_180_ (FlaB4) and Phe_182_ (FlaA2) (Fig. 2B), Val_180_ is not conserved and is located at the tip of a loop (175-184) that is shortened by 1-2 residues in the other FlaB isoforms. The second protein-protein contact, between His_208_ (FlaA2) and Thr_146_ (FlaB4), also suggests a preference for FlaB4, as in the other isoforms Thr_146_ is either absent or replaced by a lysine or alanine (Fig. 2B). Finally, for the contact between Pro_66_ of FlaA2 and Met_157_ of FlaB4, Pro_66_ is substituted in FlaB3 and FlaB2 with a glycine and an alanine respectively, likely attenuating the interaction strength (Fig. 2B). Similarly for FlaAP, of the four glycan interactions, two of these involve glycans only present in FlaB4 (Ser_182_, on two adjacent protofilaments). Thus, the majority of glycan contacts by FlaA2 or FlaAP show evidence of specificity for FlaB4.

### Purified *flaA2*^-^ flagella have a straight, not curved, morphology

A previously generated mutation of *Leptospira interrogans, flaA2*^-^, was found to be non-motile and with altered flagellar filament morphology (Lambert, Picardeau et al. 2012). Similar to *fcpA*^-^ and *fcpB*^-^ knockouts, flagella from the *flaA2*^-^ mutant were much straighter than in the wild type. However, we discovered that the coiling proteins FcpA and FcpB were both present in *flaA2*^-^ filaments, indicating that the ‘coiling’ function of FcpA and FcpB depends on FlaA2 being present.

To examine the basis of this functional behavior, we performed cryo-EM structure analysis on purified *L. interrogans flaA2*^-^ flagellar filaments. Cryo-EM micrographs revealed that these filaments were highly heterogeneous both in their diameters and curvatures (Fig. S10). A minority of the filaments (~15%) appeared nearly straight, with a diameter ~24 nm, while the majority (~83%) of the imaged filaments were curved and thinner in nature (~12 nm), likely reflecting a sheath-less FlaB core. Around 2% of the filaments were intermediate in both diameter (~20 nm) and curvature, appearing to contain a partial sheath. We analyzed the straight ~24 nm thick filaments, which provided a straightforward route to 3D structure analysis due their helically symmetric structure.

### FcpA and FcpB can form a helical lattice that completely encloses the *flaA2*^-^ filament core

A near-atomic resolution 3D structure of the *L interrogans flaA2*^-^ filament was obtained using cryoSPARC (Punjani, Rubinstein et al. 2017), applying the 11-protofilament helical symmetry common to bacterial flagella (see Methods) (Beatson, Minamino et al. 2006, Kreutzberger, Ewing et al. 2020). The resulting structure achieved an overall resolution of 2.9 Å, revealing a core as well as two distinct sheath layers, both of which completely enclose the FlaB core and follow its symmetry (Fig. 4). This high resolution allowed us to unambiguously identify the inner-most sheath layer as FcpA and the outer-most sheath layer as FcpB. The relationship of these sheath proteins with the core is similar to that observed in the wild-type filament (Gibson, Trajtenberg et al. 2020).

**Figure 4.**
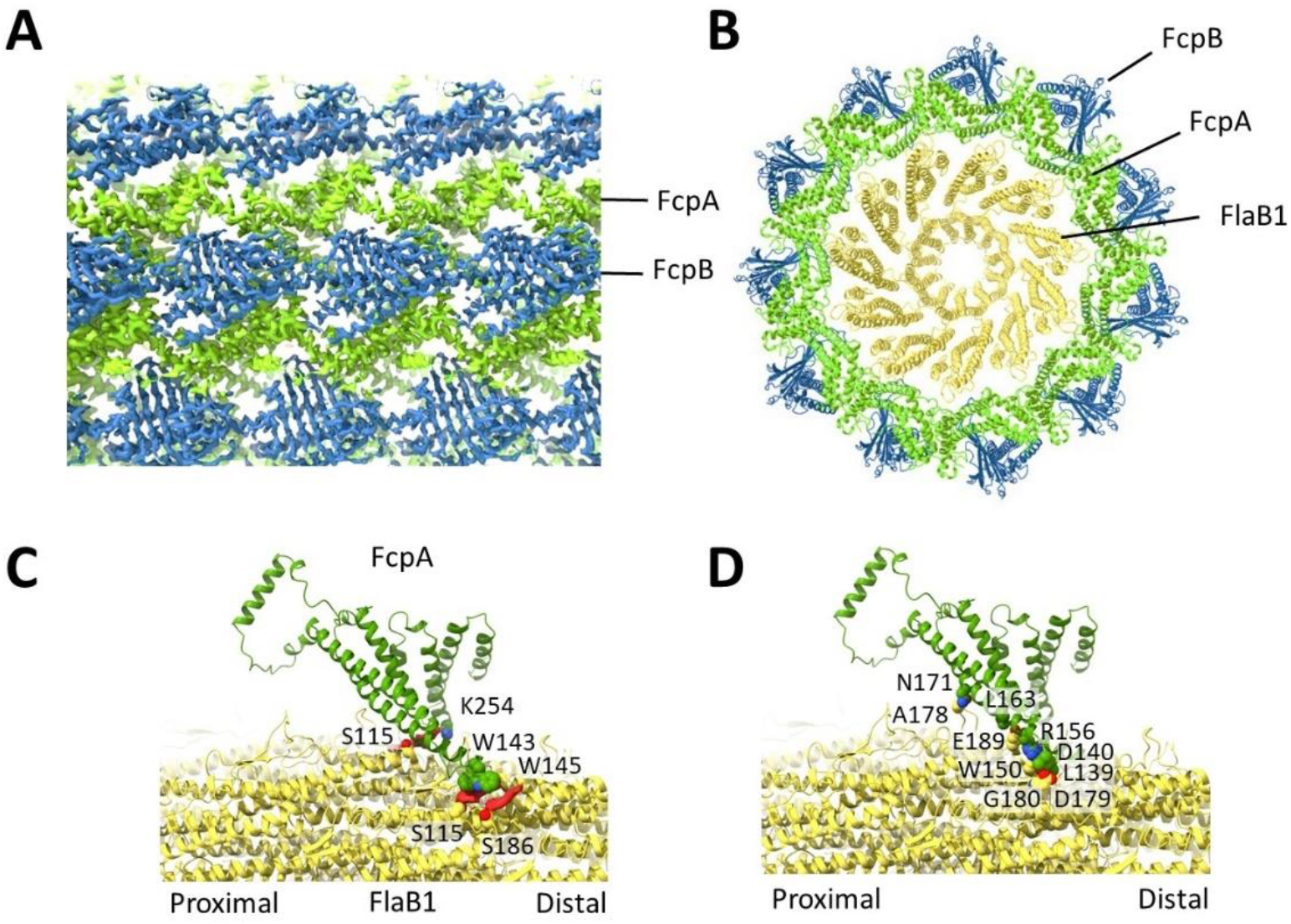
A lattice formed by FcpA and FcpB completely surrounds the core in *flaA2*^-^ filaments, and is supported by interactions with glycosylated core side chains. **A,** *flaA2*^-^ density map, highlighting the symmetric lattice of FcpA (green) and FcpB (blue) protomers. **B,** Atomic models of FlaB1 (yellow), FcpA (green), and FcpB (blue) after tracing the cryoEM maps. All three proteins conform to the 11-fold core helical symmetry, with FcpA and FcpB forming a continuous sheath that overlies the FlaB1 core. **C,** Each FcpA monomer contacts two FlaB1 protofilaments. Interactions are present between FcpA and presumably glycosylated residues of the FlaB1 core. Along one protofilament, Trp_143_ (FcpA) interacts with a glycan on Ser_115_ (FlaB1) and Trp_145_ (FcpA) interacts with a glycan on Ser_186_ (FlaB1). Lys_254_ (FcpA) interacts with a glycan on Ser_115_ of the adjacent FlaB1 protofilament. **D,** Protein-protein contacts between the FcpA sheath and the FlaB1 core occur at four locations, and mostly involve hydrophobic residues.

Each FcpA monomer is composed of 10 helices arranged in a ‘Y’, closely matching the crystal structure (San Martin, Mechaly et al. 2017, Gibson, Trajtenberg et al. 2020): one arm is formed by two long helices (α3,4), and the other arm is formed by a helical bundle (α6-10). Each FcpA contacts two adjacent FlaB protofilaments as well as six FcpA molecules (two on the same protofilament, and two on each neighboring protofilament) (Fig. 5A).

**Figure 5.**
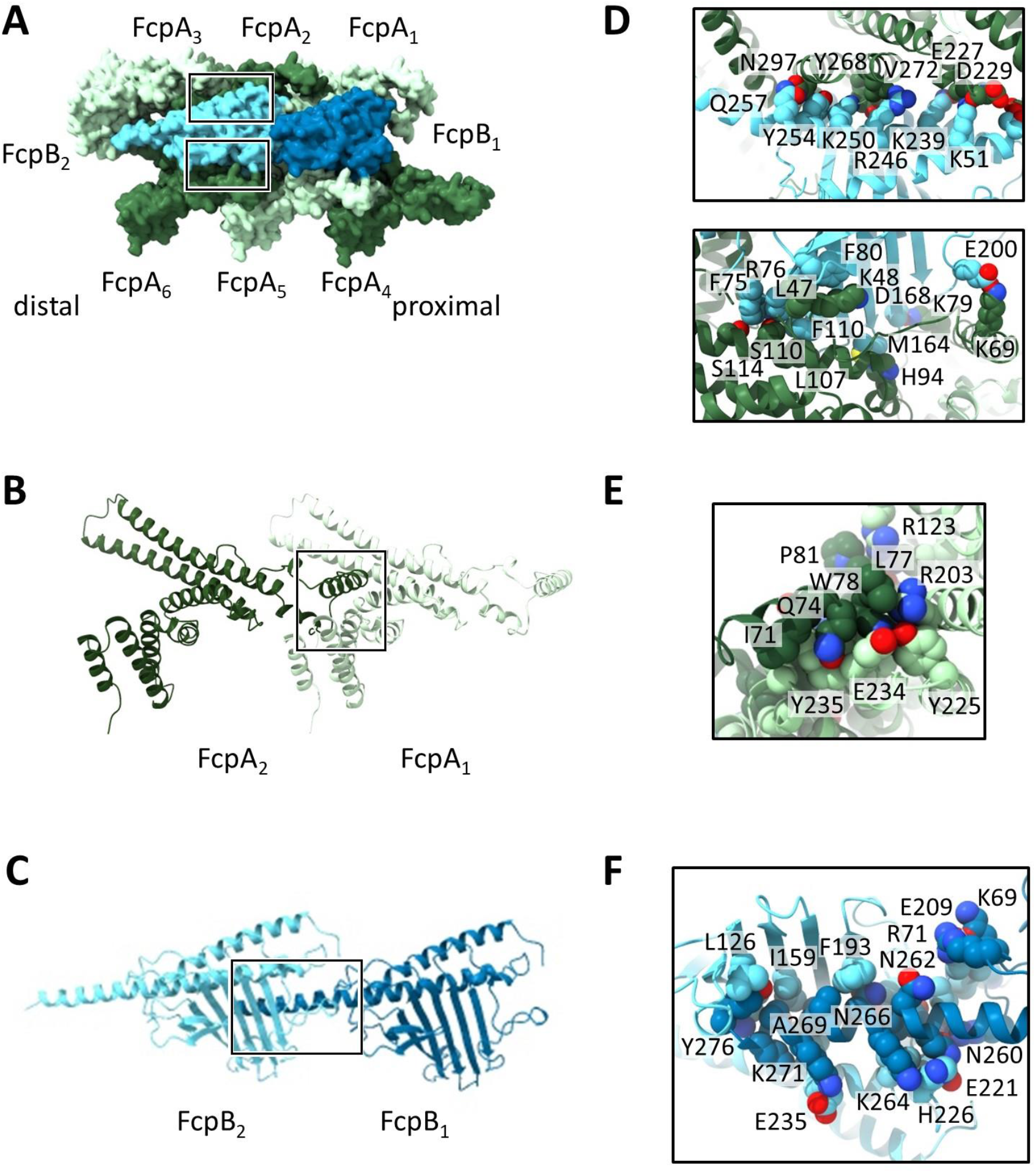
The FcpA/FcpB lattice is characterized by extensive longitudinal interactions. **A,** FcpB overlies a lattice of FcpA, with each FcpB contacting five FcpA monomers. Contacts with two lattices are shown within the boxes; top box corresponding to the top panel in **D**, and the bottom box corresponding to the bottom **D** panel. **B,** Extensive lateral FcpA interactions. The extended helix of one FcpA monomer contacts the proximal neighbor monomer. **C,** Lateral FcpB interactions occur through its long helix. **D,** FcpA and FcpB interactions. Top panel, contacts between FcpB_2_ and FcpA_2_. Bottom panel, contacts between FcpB_2_ and FcpA6. **E,** Close-up view of the axial FcpA interactions, including hydrophobic interactions with Trp_78_ of FcpA_2_ and a helix of FcpA_1_. **F,** Lateral interactions of FcpB involve contacts between the beta-sheet of one monomer and the helix of the proximal FcpB neighbor.

Numerous longitudinal interactions occur within each FcpA row. Residues 65-96 of one FcpA monomer (located on α1-2) interacts extensively with the proximal FcpA, including an especially prominent hydrophobic interaction between Trp_78_ (on the α1-2 loop) and the α7-8 segment (residues 225-240) of the next proximal FcpA monomer (Fig. 5E). This N-terminal region of FcpA within the assembly is starkly different from the crystal structure, with α1 repositioned ~52 Å in the proximal direction (Fig. S6C). This displacement removes α1 from a hydrophobic binding pocket on the same FcpA protomer, as seen in the crystal structure, and places it instead within the corresponding pocket of the next longitudinally adjacent FcpA neighbor. Similar to a domain swap, this α1 rearrangement tightly links FcpA subunits together within each individual row. Additional contacts are formed between an FcpA monomer and two FcpAs located on each longitudinally adjacent row. This network of FcpA interactions in the *flaA2*^-^ filament forms a strong, symmetric FcpA lattice around the entire FlaB core.

The outer-most sheath layer is completely accounted for by FcpB, which has two helices and a seven-stranded beta-sheet, similar to the crystal structure (Fig. S6D). FcpB forms rows which overlie two FcpA rows. FcpB protomers associate longitudinally within each row but are widely separated from adjacent FcpB rows. Longitudinal contacts are characterized by extensive interactions between a long, C-terminal helix of one FcpB (FcpB_1_) with the beta sheets and helix of the distal neighboring monomer (FcpB_2_) (Fig. 5C,F). These interactions are supported by additional contacts of extended loops from both monomers. The involvement of a long, relatively incompressible alpha helix in these extensive longitudinal FcpB interactions may impart rigidity to this region of the sheath. This implicit mechanical stability contrasts with the relatively flexible longitudinal interactions (‘tentacles’) observed for FlaA2 in our *fcpA*^-^ structure. These differences are likely important for the contrasting functions we infer for FlaA2 *vs* FcpB (see below).

There are extensive contacts between FcpA and FcpB, with each FcpB interacting with five underlying FcpA monomers (Fig. 5A). Minor contacts are made between FcpB_2_ and three of the underlying FcpAs (FcpA_1_, FcpA_3_, and FcpA_5_), while major interactions occur with FcpA_2_ and FcpA_4_ (Fig. 5D). This extensive lattice network of FcpA and FcpB aids in the formation of a stable sheath layer, with correspondingly high local resolution in this sheath region (3.0 – 3.3 Å) (Fig. S6B).

### Core interactions of FcpA and FcpB coiling proteins are mediated by FlaB1 glycosylation sites

The core region of the *flaA2*^-^ filament density closely resembles the core structure found in our *fcpA*^-^ mutant, except that it is straight and helically symmetric rather than curved (Fig. 4B). Thus, no ‘seam’ is present. Five large, globular densities, corresponding to sites of glycosylation, were seen in each FlaB monomer. The five glycosylated residues observed in the *flaA2*^-^ mutant are also the location of glycan modifications in *fcpA*^-^ flagella, although the *fcpA*^-^ filaments contained two additional sites of glycosylation (the FlaB4-specific Thr_137_ and Ser_182_).

The glycan fingerprint of the *flaA2*^-^ filaments did not allow for unambiguous FlaB isoform identification (as was possible in the *fcpA*^-^ filaments), as three of the FlaB isoforms have a Ser/Thr underlying each of the five sites of glycosylation. Instead, density at a divergent loop (residues 176-183) and at unique bulky residues in each isoform identified FlaB1 as the predominant isoform in these mutant filaments; as in the *fcpA*^-^ core, there is likely a mixture of isoforms, with the dominant isoform providing the strongest signal.

Three sheath-glycan contacts are made between each FcpA and FlaB; two of the FcpA residues are tryptophans, likely reflecting pi-stacking with the glycan sugars (Fig. 4C) (Samanta and Chakrabarti 2001). These FcpA-glycan interactions occur on both of the underlying FlaB protofilaments. One extended loop of FcpB (residues 111-124) interacts with the FlaB core, with one positively-charged FcpB residue contacting a core glycan.

Few protein-protein contacts are present between the Fcp sheath proteins and the underlying FlaB core. One contact is made between FcpB and FlaB residues, involving the same extended loop that contacts the core glycan. Five protein-protein contacts are made between FcpA and FlaB, across two FlaB protofilaments (Fig. 4D); similar to the FlaA2/FlaB4 interactions, these contacts are mostly hydrophobic in nature.

### Coiling proteins elongate the helical lattice of the filament

We measured the longitudinal repeat spacing of wild-type and mutant structures by taking the center of mass of the FlaB protomers. The average spacing for each filament remains close to constant (52.1 Å), except for the *flaA2*^-^ mutant, where it increased by ~0.5 Å (52.6 Å) (Fig. S7). Differences between the longitudinal spacings at the inner curvature and the outer curvature directly correlate with the overall filament curvature, with the largest spacing difference in the tightly coiled wild-type sample (51.1 *vs* 52.9 Å); the *fcpA*^-^ filaments are less curved, and thus have a smaller spacing difference (51.9 *vs* 52.2 Å). While the *flaA2*^-^ mutant structure is straight, the overall filament spacing is increased to a value that exceeds any part of the *fcpA*^-^ mutant structure and approaches that of the wild-type filament outer curvature. Taken together, these observations indicate that decoration by the coiling proteins in the absence of FlaAs/FlaAP, stretch and straighten the core filament.

### FlaA2/FlaAP and FcpA/FcpB segregate to opposite sides of the wild-type filament

The structure of the *Leptospira* wild-type flagellum has been challenging to solve at high resolution due to the flattened supercoiling geometry which traps filaments in a preferred orientation in the specimen ice layer(Gibson, Trajtenberg et al. 2020). Previously, we resorted to cryo-subtomogram averaging to overcome this obstacle, but the resolution of the resulting structure was highly anisotropic and limited to ~1 nanometer at best (Gibson, Trajtenberg et al. 2020). However, we serendipitously discovered spontaneous filament fragmentation in aged wild-type specimens. The resultant shorter pieces more freely reoriented in the cryo-EM grids (see Methods). By collecting image data sets of 45°-tilted samples, we were therefore able to collect sufficient views to solve an isotropic reconstruction of the wild-type filament using single-particle cryo-EM methods (Fig. 6; see Methods).

**Figure 6.**
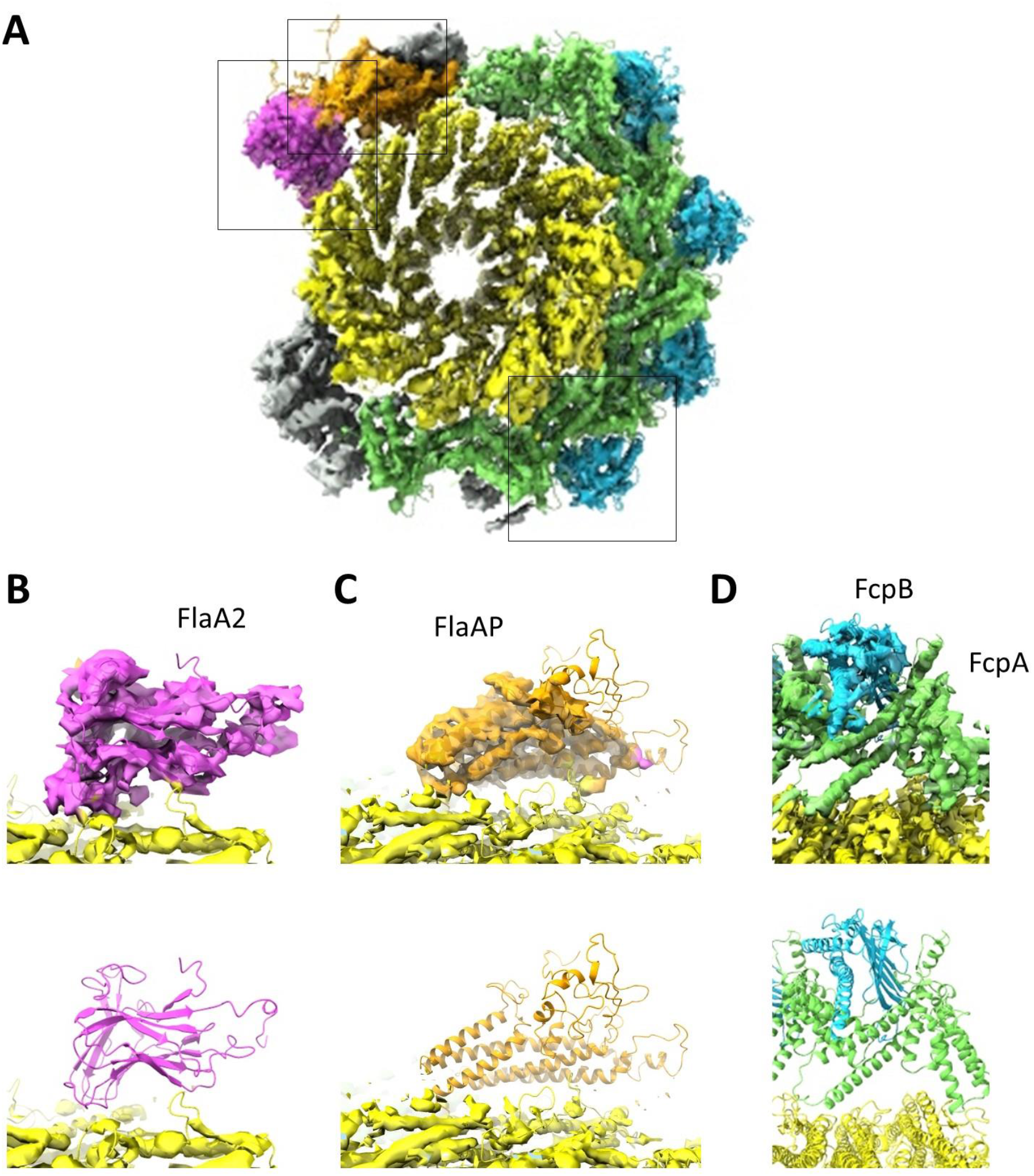
Lattice of FcpA and FcpB coiling proteins is interrupted by FlaA2/FlaAP in the wild-type *Leptospira* filament. **A,** Cross-sectional view of the wild-type filament structure. Density corresponding to the core is colored yellow, to FlaA2 is pink, to FlaAP is orange, to FcpA is green, and to FcpB is blue. Additional sheath density that does not correspond to these proteins is in gray. The three boxes highlight the regions featured in B, C, and D. **B,** Zoomed-in view of FlaA2 fit into the wild-type density. The top panel shows the density, the bottom shows the fit of the model within the density. FlaA2 is colored pink, and the FlaB model is colored yellow. **C,** FlaAP (orange) fit into the density, as in B. **D,** The fit of FcpA (green) and FcpB (blue) into the density, as in B.

The resolution achieved in the 3D reconstruction (6.5 Å in the best resolved regions) allowed unambiguous fitting of FlaB, FcpA, FcpB, FlaA2, and FlaAP. The overall architecture of the core and sheath proteins is consistent with the previously reported structure obtained by cryo subtomogram averaging (Gibson, Trajtenberg et al. 2020). Moreover, the detailed structure and interfaces of the coiling proteins FcpA and FcpB are consistent with our new high-resolution *flaA2*^-^ mutant structure; similarly, the detailed structure interfaces of FlaA2 and FlaAP are consistent with our high-resolution *fcpA*^-^ structure.

In contrast to the *flaA2*^-^ structure, however, in the wild-type structure the FcpA/FcpB helical lattice is incomplete and asymmetric. These coiling proteins are completely displaced in 5 protofilaments by FlaA2, FlaAP, as well as several additional globular densities, which likely correspond to extra sheath proteins like FlaA1 and others not yet identified. The position of FlaA2 and FlaAP on the inner curvature of the wild-type filament approximately matches that found in our *fcpA*^-^ structure. In sum, comparison of our new wild-type and mutant structures indicates that FlaA2, FlaAP, and additional unidentified components displace part of the FcpA/FcpB helical lattice in the wild-type structure (gray regions in Fig. 6).

## Discussion

Here we have presented three *Leptospira* filament structures: one from wild-type flagella, and two from specific sheath protein KO mutants (*fcpA*^-^ and *flaA2^-^*). The structures show evidence of FlaB glycosylation, which appears to be crucial for sheath protein binding. When the Fcp coiling factors are missing (as in the *fcpA*^-^ mutant), the sheath proteins bind only to the inner curvature of the core; in contrast, when the FlaA2 sheath factor is missing (as in the *flaA2*^-^ mutant), the coiling factors bind symmetrically around the entire core. This supports a role of FlaA2 as a ‘templating’ factor, preventing the Fcp sheath proteins from binding along the inner curvature and segregating the ‘templating’ and ‘coiling’ factors to opposing sides of the wild-type filament (Fig. 7).

**Figure 7.**
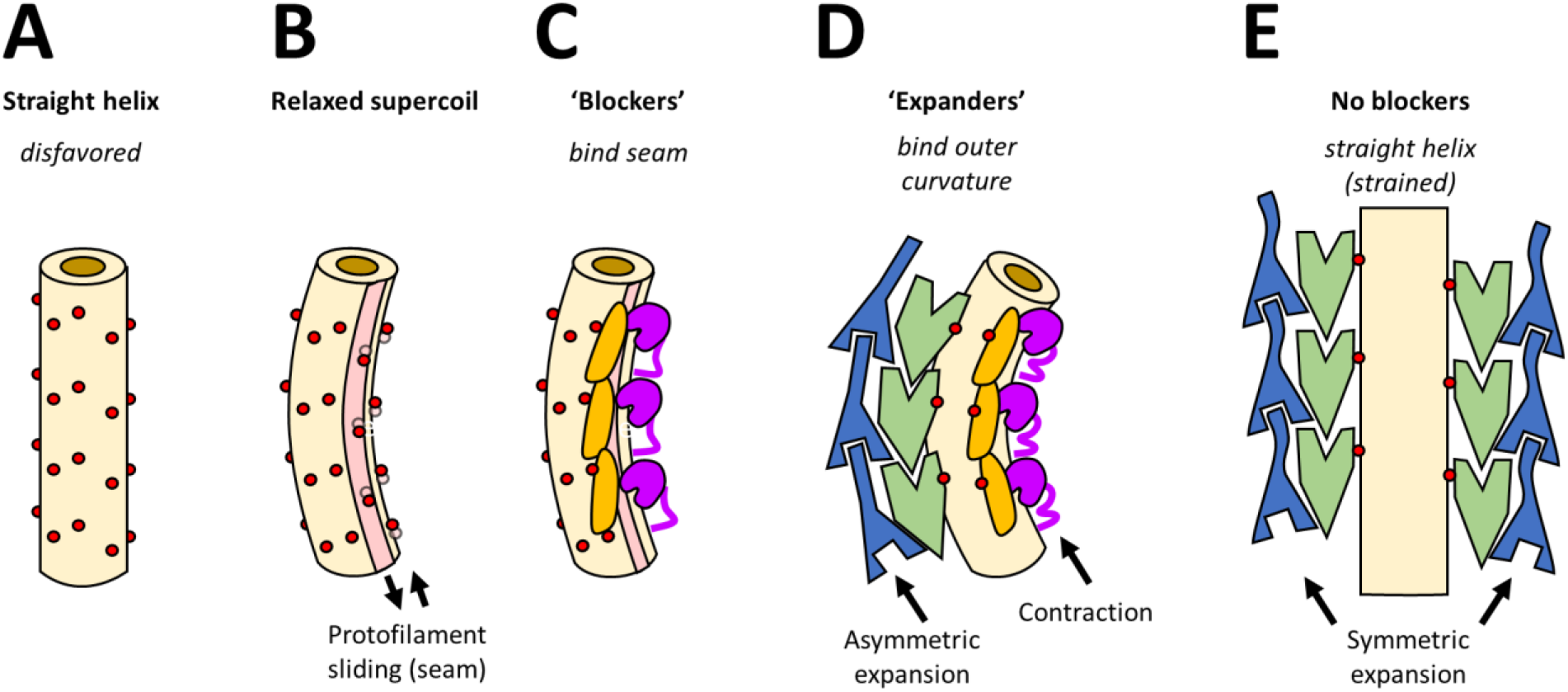
Model for joint cooperative activation of *Leptospira* flagellar filament function by multiple sheath proteins. **A,** Straight, helically symmetric core conformation; note that this is expected to be energetically disfavored. **B,** In the absence of sheath proteins, the FlaB core (yellow) assumes a relaxed supercoiled shape, like the ‘normal’ form observed in *Salmonella* and other bacteria. Red dots denote glycosylated core surface residues, following the filament helical lattice. A seam (pink) associated with the relaxed supercoil would feature sliding of several neighboring protofilaments, disturbing the helical lattice (**A**) on the inner curvature side of the filament. Specific targeting of FlaA2 (magenta) and FlaAP (orange) to the seam would then block binding of additional sheath proteins on the inner curvature (**C**). FcpA (green) and FcpB (blue) then function as ‘expanders’ upon binding the core (**D**), pushing against their longitudinal neighbors via specific binding interactions involving rigid secondary structure elements. Localization of FcpA/FcpB to the outer curvature (due to blocking by FlaA2, FlaAP and potentially other sheath proteins) thus promotes asymmetric lattice expansion on the outer curvature side of the filament. The fully assembled filament would thus adopt a tightly coiled form, enabling motility. Our wild-type structure indicates that several additional sheath proteins bind the inner curvature side of the filament (not shown). However, if FlaA2 and FlaAP are not present (as in the *flaA2*^-^ mutant), loss of blocking function enables the FcpA/FcpB lattice to displace all other sheath components and fully envelop the FlaB core. The resulting filaments are helically symmetric and straight (**E**), and hence unable to support motility.

### Polymorphic switching in the supercoiled filament

The curved nature of the *fcpA*^-^ filaments suggests that each FlaB protofilament will have variations from its neighbor. However, we found that one protofilament (#4) exhibited a stark difference, with a ~2 Å lateral shift in the D1 domain relative to the other ten protofilaments (Fig. 7B). These other protofilaments resemble ‘L’-type conformations observed previously in other bacteria, while the shifted protofilament resembles the ‘R’-type (Wang, Burrage et al. 2017). These observations suggest that the *fcpA*^-^ mutant supercoiled form may contain a 10L/1R composition. The FlaA2 and FlaAP sheath factors are found along this ‘R’ protofilament, suggesting that this ‘R’ transition may be crucial for FlaA recruitment and binding (Fig. 7C). Further work will be required to determine the polymorphic form of the wild-type core, as it is probable that it too may contain a mixture of ‘L’ and ‘R’ states.

### Distinct functional roles of ‘templating’ versus ‘coiling’ sheath proteins

The FlaA sheath factors appear to ‘template’ the core, preventing the binding of the remaining sheath proteins to the occupied inner curvature. The presence of these localized proteins ensures that the filaments will adopt a curved form, as either a ‘relaxed’ supercoil (when the Fcp proteins are absent; Fig. 7C) or a tight supercoil (when the Fcp proteins are also present; Fig. 7D). This role suggests that these FlaA proteins are recruited to the growing filament before the Fcp proteins, allowing them to bind along the inner curvature of the otherwise unoccupied FlaB core.

However, when the FlaA proteins are absent, FcpA and FcpB coiling factors are able to bind symmetrically around the filament, forcing the filament into a straight form (Fig. 7E). By ‘stretching’ the lattice, these coiling proteins thereby increase the lattice spacing of the core in both the *flaA2*^-^ filaments and along the outer curvature of the wild-type filaments. Stretching along the outside of the wild-type filament would induce the formation of tight coils associated with motility. FcpA and FcpB coiling factors have not been identified in other spirochetes and may be unique to *Leptospira* (Wunder, Figueira et al. 2016, Wunder, Slamti et al. 2018). This observation correlates with the fact that purified flagella from other spirochetes are curved, but do not form tight supercoils. It thus remains unclear what function(s) FlaA may perform in other spirochetes, despite its widespread conservation across this phylum.

### Role of glycosylation in sheath factor recruitment

We were able to structurally identify seven key glycosylation sites within the FlaB core of the *fcpA*^-^ filament, four of which appeared crucial for interactions with the FlaA2 sheath, three which appeared crucial for interactions with FlaAP, and two that did not appear to influence sheath binding. Of the five glycans in the *flaA2*^-^ structure, three are involved in sheath binding (two for FcpA, and one for FcpB). It is probable that the core of the wild-type filament is similarly glycosylated, though these features cannot be resolved with the current structure. Three *Leptospira* glycosylation sites (Ser_115_, Ser_126_, Thr_137_) were also identified as locations of glycosylation in the FlaB core of *T. denticola* (Kurniyati, Kelly et al. 2017); while Ser_115_ and Ser_126_ are present in all *Leptospira* FlaB isoforms, Thr_137_ is only amenable to glycosylation in FlaB4, and is not present in our *flaA2*^-^ structure. In both *Leptospira* and *T. denticola*, two of these glycosylation sites lie within the consensus sequence predicted to bind the flagellin-recognizing toll-like receptor 5 (TLR5), and glycosylation therefore may interfere with its ability to recognize the flagellum (Kurniyati, Kelly et al. 2017, Holzapfel, Bonhomme et al. 2020), though the overall periplasmic location and the sheathed character of the filament must also contribute to a lack of recognition by TLR5 (Holzapfel, Bonhomme et al. 2020). Many of these *Leptospira* glycosylation sites are conserved amongst spirochete FlaBs (Kurniyati, Kelly et al. 2017), suggesting that similar modifications may be observed in the FlaB core of other species, providing roles in sheath-protein binding, and evasion from the host immune response.

### Role of FlaB isoforms

Based on the glycosylation patterns and mass spectrometry analyses, we identified the core of the *L. biflexa fcpA*^-^ filament as being composed primarily of FlaB4. Previous studies of wild-type *L. interrogans* showed a vast discrepancy in the number of copies of each FlaB isoform, ranging from ~12,000 copies per cell of FlaB1 to ~300 copies per cell of FlaB3 (Beck, Malmstrom et al. 2009). Therefore, it was surprising to identify FlaB4, and not FlaB1, as the major component of all 11 protofilaments of our *fcpA*^-^ structure, even though both isoforms are equally abundant in our *fcpA*^-^ filament samples (Table S1). On the one hand this could reflect genuine differences in protein stoichiometries between *L. interrogans* and *L. biflexa* flagella. On the other hand, perhaps other regions of the core (with no FlaA2/FlaAP bound) are more heterogeneous in terms of FlaB composition. As FlaB4-specific glycosylation sites and loops appear critical to the interaction between the sheath and the core, it is likely that this isoform is required for FlaA2 and FlaAP binding. Whether this isoform is dominant in the wild-type *L. biflexa* structure remains unknown, as the resolution of the core in our current wild-type structure unfortunately barred a detailed analysis.

In the *L. interrogans flaA2*^-^ filament, however, the major isoform appeared to be FlaB1. The sheath-glycan interactions and most of the sheath-core protein-protein contacts are not FlaB1-specific, raising the possibility that the Fcp sheath factors may have the ability to bind to various FlaB isoforms. This could explain the ability of these coiling factors to assemble symmetrically in these mutant filaments, as these Fcp sheath factors would still be able to bind even if the inner curvature of the core consisted of a different FlaB isoform.

### Role(s) of additional, undefined sheath proteins

While mass spectrometry analyses of the purified flagella indicate that FlaA1 is present in the *fcpA*^-^ filaments (Table S1), we were unable to identify it in our structure, suggesting that FlaA1 is not as stably bound to the core. A previous study of *L. biflexa* failed to detect FlaA1 in purified *fcpA*^-^ filaments, though the sheath factor was present in the cell lysate (Sasaki, Kawamoto et al. 2018); this contrasts results from *L. interrogans* in which FlaA1 was indeed detected in purified *fcpA*^-^ filaments (Wunder, Figueira et al. 2016). The presence of additional decorated protofilaments in a minority of our analyzed *fcpA*^-^ filaments may reflect the presence of this and/or other sheath components.

The wild-type structure appears to contain additional sheath factors, with densities that do not correspond to known components −FlaA2, FlaAP, FcpA, FcpB, or the FlaB core (Fig. 6A). While one of these densities is likely FlaA1, the fitting of a predicted model was not conclusive, due to limited local resolution of the map segments, and/or actual model inaccuracies. Additional proteins are anticipated, which would correspond to novel flagellar protein species, future studies shall shed light into their identities, precise location and roles in endoflagellar filament assembly. Altogether, these additional factors are also located near the inner curvature of the filament, adjacent to the FcpA coiling factors. These factors may therefore also play a role in ‘templating’ the sheath, likely stabilizing the wild-type asymmetric sheath arrangement or otherwise facilitating its formation.

### Conclusion

The *Leptospira* flagellum is a complex assembly, composed of multiple different types of proteins- a glycosylated FlaB core, FlaA ‘templating’ factors, and Fcp ‘coiling’ factors. The simultaneous localization of the FlaA sheath factors to the inner curvature and the Fcp factors to the outer curvature, results in a functional, supercoiled filament; without both types of sheath proteins, the filament remains relaxed and is unable to bend into the tight coils that are associated with motility. Continued investigations of spirochete flagellar architecture will help to inform how these remarkable filaments play such a vital role in the unique movement of this important bacterial phylum.

## Methods

### Strains and culturing of *Leptospira*

Cultures of *Leptospira biflexa* serovar Patoc strain Patoc I (Paris) wild-type and *fcpA*^-^ mutant samples (Wunder, Slamti et al. 2018), as well as *L. interrogans* serovar Manilae *flaA2*^-^ (Lambert, Picardeau et al. 2012), were grown in Ellinghausen-McCullough-Johnson-Harris (EMJH) liquid medium at 30°C (Wunder, Figueira et al. 2016).

### Flagella purification

Purification of *L. biflexa fcpA*^-^ and *L. interrogans flaA2*^-^ flagella was performed as previously described (Miller, Miller et al. 2016). Briefly, 500mL of a *fcpA*^-^ or *flaA2*^-^ culture was harvested and centrifuged at 8000xg for 15 minutes, and the pellet was washed with cold PBS and re-centrifuged. The pellet was resuspended in 30mL Tris buffer (150mM Tris (hydroxymethyl amino methane, pH 6.8, with 0.9% sodium chloride) and centrifuged as before. The pellet was then resuspended in 15mL of Tris buffer, and stirred at 4°C for 10 minutes, before 1.5mL of 20% TritonX-100 was added to the sample. After stirring at room temperature for 1 hour, the sample was centrifuged at 15,000xg for 45 minutes, and the pellet was resuspended in 15mL Tris buffer. 1000 units of mutanolysin was added dropwise to the sample, which was stirred at room temperature for 1 hour, and then overnight at 4°C. The sample was then centrifuged at 8,000xg for 30 minutes. The pellet was discarded, and 2.2mL of ammonium sulfate was added to the supernatant (for a final ammonium sulfate concentration of 12.6%); this was stirred at 4°C for 20 minutes. This sample was then centrifuged at 120,000xg for 2 hours. The resulting pellet was resuspended in water, and the centrifugation was repeated. The final pellet was resuspended in 600 μL water. The sample was then analyzed with SDS-PAGE and Western blots to ensure that the expected flagellar proteins were present.

*L. biflexa* WT flagella were purified as previously described (Wunder, Figueira et al. 2016). Briefly, 500 mL of a *L. biflexa* WT culture were harvested and centrifuged at 8000xg for 15 minutes. The pellet was washed with cold PBS and re-centrifuged. The pellet was then suspended in 30mL 0.15M Tris.HCl pH8.0, 0.5M sucrose and centrifuged as before. The pellet was then resuspended in 30mL of Tris-sucrose buffer (0.5 M sucrose, Tris 150mM, 50mM NaCl), stirred at 4°C for 10 minutes, and then 3 mL 10% TritonX-100 were added. After stirring at room temperature for 30 minutes, 0.1 mg/mL (final concentration) hen egg-white lysozyme, 0.005 mg/mL DNAse, 0.01 mg/mL RNAse and 2 mM MgCl_2_ were sequentially added dropwise, further stirring at room temperature for 2 hours. The samples were then stirred 10 minutes with 2 mM MgSO_4_, and then 10 minutes with 2 mM EDTA pH8.0, then centrifuged at 17000xg for 15 minutes. The pellet was discarded, and 4 mL of 20% PEG8000, 1M NaCl were added to the supernatant and incubated for 30 min at 4°C. This sample was centrifuged at 27000xg for 30 min, and the pellet resuspended in 4 mL 150 mM Tris pH8.0, 50 mM NaCl, and slowly stirred at 4°C. Flagellar filaments were recovered by ultracentrifugation at 80000xg for 45 minutes, this final pellet was resuspended in 500 μL of 150 mM Tris pH 8.0, 50 mM NaCl.

### Cryo-EM sample preparation

3-4 μL of purified *L. biflexa fcpA*^-^ flagella samples were applied to Quantifoil R1.2/1.3 Cu 300 mesh grids (Ted Pella, Inc., Redding, CA), 3-4 μL of purified *L. interrogans flaA2*^-^ flagella samples were applied to Quantifoil R1.2/1.3 Cu200 mesh grids (Ted Pella, Inc., Redding, CA), and 2.5 μL of *L. biflexa* wild-type purified flagella were applied to Quantifoil 2.3/1.3 Cu300 mesh grids (Ted Pella, Inc., Redding, CA). All grids were plasma discharged in a Gatan Model 950 Solarus Advanced Plasma System with H2/O2 for either 30 seconds (*fcpA*^-^ and *flaA2^-^*) or for 20 seconds (wild-type). The grids were incubated for 1 minute, and then plunge frozen with a Vitrobot Mark IV, with a blotting time of 6 seconds and a blotting force of 2, all at 18 - 22°C and 100% humidity.

### Data Collection

Initial micrographs of *fcpA*^-^ filaments were collected through the program SerialEM (Mastronarde 2005) on the 200kV Thermo Scientific Glacios containing a K2 detector. Images were collected with a pixel size of 0.896 Å.

Wild-type, *flaA2*^-^, and additional *fcpA*^-^ micrographs were collected on the 300kV Titan Krios microscope, containing a K3 detector, through the program SerialEM. *fcpA*^-^ and *flaA2*^-^ images were collected with super-resolution pixel size of 0.534 Å and a magnification of 81000x. A magnification of 130000x was used for the wild-type sample, with a super-resolution pixel size of 0.525 Å. A defocus between −1.5 μm to −3.2 μm was used for the *fcpA*^-^ samples, and a defocus of −1.5 – −2.6 μm was used for the *flaA2*^-^ samples, and a defocus of −3.0 μm was used for the wild-type sample. A total dose of 60 e^-^ /Å^2^ was used for both *fcpA*^-^ and *flaA2*^-^ samples, and a total dose of 54 e^-^/Å^2^ was used for the wild-type sample. 11906 *fcpA*^-^ micrographs were collected; for 4994 of those micrographs, image shift was used to take images at four holes per stage position. 2465 *flaA2*^-^ micrographs were obtained, and 718 wild-type micrographs were collected (197 at 0° tilt, and 521 at 45° tilt). For all micrographs, only one image was taken per hole.

### RELION reconstruction of the *fcpA*^-^ filaments

#### Initial Glacios reconstruction

An initial helically symmetric model was generated from 802 micrographs collected on the Glacios. MotionCor2 (Zheng, Palovcak et al. 2017) was used for motion correction, Gctf (Zhang 2016) in Relion 3.0 (Zivanov, Nakane et al. 2018) was used for CTF correction. Manual selection of filaments was performed in EMAN (Ludtke, Baldwin et al. 1999), resulting in 74,369 image segments. 2D classification was performed to separate sheathed and bare filaments; 26,552 image segments (corresponding to filaments without a visible sheath) were selected for further reconstruction of the core. A 30Å low-pass filter of a *Bacillus subtilis* flagellar filament (EMDB-8852) was used as an initial reference (Wang, Burrage et al. 2017) for helical reconstruction. The following helical parameters were used: a 0.5° local searches were used, 11 asymmetrical units, an initial helical rise of 4.72 Å (with a 0.2 Å search between 4.42 Å and 5.02 Å), an initial helical twist of 65.3° (with a 0.2° search between 62.3° and 68.3°), a central Z length of 30%, a range factor of local averaging of 2, a psi angular search range of 10°, a tilt angular search range of 15°, an outer tube diameter of 280 Å, an initial angular sampling of 0.9°, and using fixed tilt-prior angles.

#### Initial Krios reconstruction

11906 micrographs were collected on the Krios. The initial 4461 micrographs were also processed with Relion 3.0 (Zivanov, Nakane et al. 2018). Motion correction and CTF correction were performed with MotionCor2 (Zheng, Palovcak et al. 2017) and Gctf (Zhang 2016), respectively. Filament selection was performed with crYOLO (Wagner and Raunser 2020), using filaments manually selected from 20 random micrographs with EMAN2 (Tang, Peng et al. 2007) for training, and was run with a box size of 320 pixels and a box distance of 56 pixels. This procedure yielded 807,689 image segments. Mis-selected particles were removed with 2D classification, yielding 803,924 image segments. The *pf_smooth* program (Debs, Cha et al. 2020) was used to remove discontinuities from the filament trajectories, with the following filament parameters: a rise per subunit of 4.72 Å, a twist per subunit of 65.356°, 11 protofilaments, a window size of 7, a fit order of 2, a minimum filament length of 10, a direction tolerance of 20, and a phi, theta, psi, distance, and twist tolerance of 10. After four iterations of smoothing, 430,755 image segments remained (53.3% of the crYOLO-selected image segments).

3D helical refinement was performed on these smoothed image segments. A 10 Å low-pass filtered map of the FlaB core structure from the Glacios data as a reference, and the same helical parameters as the Glacios refinement were used. To account for the asymmetry of the sheathed filament, the resulting star file was then expanded 11-fold with relion_particle_symmetry_expand (utilizing the helix function, with 11 protofilaments, a twist of 65.4°, and a rise of 4.72 Å), resulting in 4,738,305 image segments.

Due to the asymmetry and heterogeneity present in the sample, additional image analysis steps were utilized to obtain meaningful 3D reconstructions (Mentes, Huehn et al. 2018). Particle subtraction was performed prior to focused 3D classification targeting the sheath. A cylindrical mask, 90 Å in diameter and along the entire length of the reconstructed filament, was used for particle subtraction. An initial 3D classification was performed on a random subset of ~70,000 subtracted particles, using the following parameters: a 15 Å filtered copy of the helical refinement as a reference, the same mask that was used in the subtraction, 5 classes, a regularization parameter of 100, no image alignment, and no helical reconstruction. The classification converged by 27 iterations, and resulted in a sheathed class (9,942 particles, or 14.2% of asymmetric subunits), three bare classes (totaling 48,371 particles, or 69.0% of asymmetric subunits), and a class possibly corresponding to a part of the sheath that was cut off by the mask (11,823 particles, or 16.9% of asymmetric subunits). These classes were then used as a seed for the subsequent 3D classification on the full ~4 million particle dataset, utilizing the same parameters for classification. This run converged after 16 iterations, and resulted in one decorated sheath class (416,720 particles, or 8.8% of asymmetric subunits), three bare classes (totaling 4,011,067 particles, or 84.7% of asymmetric subunits), and a final class that may represent part of the sheath that was cut off by the mask (310,518 particles, or 6.6% of asymmetric subunits).

#### Final Krios reconstruction

All 11906 micrographs were then processed using Relion3.1 (Zivanov, Nakane et al. 2020). Motion correction, defocus estimation, and particle selection were performed as before, resulting in 1,632,891 total image segments. Roughly half of these image segments did not have converged alignment parameters, with numerous discontinuities in the filament trajectories, hindering initial efforts at helical processing. An initial alignment was therefore generated using one round of Relion refinement (Class3D) with a fine-grained, exhaustive search. The following parameters were used: a regularization parameter of 4, an angular sampling interval of 1.8°, an offset search range of 20 pixels with a 1 pixel search step, with no local angular searches performed, and using helical reconstruction with a tube outer diameter of 170 Å, a tilt search range of 15°, a psi search range of 20°, with fixed tilt-priors, and without applying helical symmetry. The reference for this step was the sheathed class from the original 4461-micrograph analysis. Utilization of an 80 Å high-pass filtered reference improved the ability to track continuously along filaments, possibly by de-emphasizing sheath features that could interfere with the alignment. After this classification, *pf_smooth* was applied using the same parameters as before, resulting in 1,538,897 image segments. 2D classification was used to discard ~1000 mis-selected image segments, and Relion 3D helical refinement was performed on the remaining 1,538,130 image segments, using the same parameters as the initial Krios reconstruction. Particle expansion (yielding 16,919,430 image segments) and subtraction were performed as before. Asymmetric 3D classification was then performed, seeded by the classes identified in the 4461-micrograph analysis, with convergence achieved after 19 iterations. This resulted in a decorated sheath class (816,888 particles, or 4.8% of asymmetric subunits), three bare classes (14,276,994 particles for 84.4% total of asymmetric subunits), and a final class that may represent part of the sheath cut-off by the mask (1,825,548 particles, or 10.8% of asymmetric subunits). An additional round of focused classification was performed on the sheathed class, utilizing a smaller cylindrical mask focused on the FlaA2 density. This converged after 38 iterations, resulting in one class with strong density (300,616 particles, corresponding to 36.8% of the particles in the second classification and 1.8% of the overall particles), one class with moderate density (40,595 particles, corresponding to 5% of the particles in the second classification and 0.2% of particles overall), and two classes with poorly defined density (475,677 particles corresponding to 58.3% of the second classification and 2.8% of all the particles).

### Reconstruction of *flaA2*^-^ filaments

The *flaA2*^-^ micrographs were initially processed with Relion3.1 (Zivanov, Nakane et al. 2020), with motion correction and CTF correction performed by MotionCor (Zheng, Palovcak et al. 2017) and gctf (Zhang 2016), respectively. Filaments were selected with crYOLO (Wagner, Lusnig et al. 2020); the program was trained using filaments manually selected from 20 micrographs with EMAN2 (Tang, Peng et al. 2007). All flagellar filaments (including those skinny and thick, straight and curved) were selected; resulting in 251,215 particles from 9670 filaments. The crYOLO filament selections were manually separated into a straight/thick, intermediate/curved, or skinny/curved group. The straight, thick filaments (~24 nm in diameter) were used for all subsequent analysis, consisting of 36,151 image segments. These filaments were extracted in Relion with a box size of 384, and an initial 3D volume was generated using 11-fold symmetry, using a low-passed volume of the *fcpA*^-^ core model as a reference.

All subsequent refinement steps were carried out in cryoSPARC (Punjani, Rubinstein et al. 2017). To ensure that the majority of the filaments were thicker and straight, one round of 2D classification was performed; no particles were discarded. Helical refinement was then performed, using a low-passed volume of the Relion-generated structure as an initial reference. For all steps, 11-fold symmetry, with a twist of 65.3° and a rise of 4.72 Å, was used. Global CTF refinement and local CTF refinement were performed, before an additional round of helical refinement, utilizing the same parameters as before. A resolution of 2.9 Å was reported for the final helical reconstruction.

### Reconstruction of wild-type filaments

Gain referencing was performed in IMOD (Kremer, Mastronarde et al. 1996) using the program ‘clip’, and the particles to a pixel size of 2.19 Å using the IMOD program ‘newstack’. All further processing and refinement steps were carried out in cryoSPARC v.3.3.1 (Punjani, Rubinstein et al. 2017). The Patch Correction tool (Rubinstein and Brubaker 2015) was used for motion correction, followed by multi-frame patch CTF estimation; default parameters were used for each. The filament tracer tool (in template-free mode) was used for filament selection, using a 300 Å diameter and a fractional separation distance of 0.173. 40,061 particles were extracted, with a box size of 192 pixels. Per-particle local motion correction was applied, and misselected particles were removed with 2D classification; this left 36,638 particles. Helical refinement was performed with the following parameters: an initial mask based on the tomographic wild-type structure (Gibson, Trajtenberg et al. 2020), automasking enabled, and using initial helical parameters of a 0° twist, a 52 Å shift, and a maximum initial tilt search of ±20°. The resultant structure had a reported resolution of 5.44 Å. Local per-particle CTF correction was performed (using default parameters) before repeating helical refinement; this structure gave a reported resolution of 4.5 Å. Global CTF refinement (including beam tilt, spherical aberration, and higher order trefoil and tetrafoil terms) and local CTF refinement were repeated once, followed by an additional round of helical refinement. The resulting structure reported a resolution of 4.28 Å, roughly the Nyquist limit of the sample. Heterogeneous refinement was used, utilizing two classes based on sheath density. The class with a more complete sheath contained 26,788 particles. Additional rounds of helical and non-uniform refinement further improved the high-frequency signal.

### Mass Spectrometry

#### Flagella purification for LC-MS/MS

*L. biflexa fcpA*^-^ were purified for mass spectrometry using the method described above for the wild-type samples. Protein concentration of the extracts were determined by SDS-PAGE and densitometry analysis using the LMW-SDS Marker Kit (GE Healthcare) as standard.

#### Nano LC-MS/MS analysis and protein identification

Flagellar extracts were run on 12% acrylamide SDS polyacrylamide gels and processed as previously described (Rossello, Lima et al. 2017). Briefly, 20 μg of proteins of each replicate were run until samples entered 1 cm into the resolving SDS-PAGE. After slicing, each band was destained and cysteine alkylation was performed in-gel by successive incubation with 10 mM dithiothreitol for 1 h at 56 °C and then 55 mM iodoacetamide at room temperature for 45 minutes. In-gel proteolytic digestion was performed overnight at 37°C using sequencing-grade trypsin (Promega). The resulting peptides were extracted at room temperature with 50% acetonitrile/0.1% trifluoroacetic acid. Peptides were desalted using C18 microcolumns (ZipTip^®^ C18, Millipore), vacuum dried and resuspended in 20 μL of 0.1% formic acid.

Three biological replicates of flagella purifications were analyzed using a nano-HPLC (UltiMate 3000, Thermo) coupled to a Q-Orbitrap mass spectrometer (Q Exactive Plus, Thermo). Tryptic peptides (5 μg) were separated into a 75 μm × 50 cm, PepMapTM RSLC C18 analytical column (2 μm particle size, 100 Å, Thermo) at a flow rate of 200 nL/min using a 90 minutes gradient (from 1% to 35% of acetonitrile in 0.1% formic acid). Mass analysis was performed in a data-dependent mode (full scan followed by MS/MS of the top 12 m/z in each segment) using a dynamic exclusion list.

PatternLab for Proteomics (Version V, http://www.patternlabforproteomics.org/) was used for protein identifications (Carvalho, Lima et al. 2016). Briefly, raw data were searched against a target decoy database including *Leptospira biflexa* serovar Patoc_UP000001847 sequences downloaded from Uniprot (December, 2021) and 127 most common mass spectrometry contaminants. Search parameters were set as follows: enzyme: trypsin; enzyme specificity: full specific; oxidation of methionine as variable modification and carbamidomethylation as fixed modification; 35 ppm of tolerance from the measured precursor m/z. Peptide spectrum matches were filtered using the Search Engine Processor (SEPro) using the following parameters: acceptable FDR: 1% at the protein level; a minimum of two peptides per protein and 10 ppm precursor mass tolerance. Patternlab for proteomics’s Venn Diagram module was used to identify proteins present in all replicates. The mass spectrometry proteomics data have been deposited to the ProteomeXchange Consortium via the PRIDE (Perez-Riverol, Csordas et al. 2019) partner repository with the dataset identifier PXD030741.

### Model building

AlphaFold2 (Jumper, Evans et al. 2021) was used to generate structural predictions for the *L. biflexa* proteins FlaB1, FlaB2, FlaB3, FlaB4, FlaA2, and FlaA1. For the *fcpA*^-^ core region, each FlaB AlphaFold2 model was manually fit into the density in ChimeraX (Goddard, Huang et al. 2018), and then Isolde (Croll 2018) was used to individually improve the fit of each isoform (Table S2). We identified FlaB4 as the presumptive isoform present in the sample (based on the glycan fingerprint), and used Isolde to individually fit the FlaB4 monomer into each of the 11 unique protofilaments.

In the *fcpA*^-^ sheath region, the AlphaFold2 FlaA2 model was manually aligned to the beta-sheet density in ChimeraX, and then Isolde was used to fit the sequence into the density. Bulky side chains were fit into lobes in the corresponding density, helping to confirm the sequence registration and helping to confirm the identity of the density as FlaA2.

Structural predictions of uncharacterized proteins identified in abundance in our sample through mass spectrometry were also obtained with AlphaFold2. One protein, the product of LEPBI_I0551, was predicted to form an alpha-helical bundle, similar to the observed helical sheath density associated to FlaA2. This model was manually aligned into our density with ChimeraX (Goddard, Huang et al. 2018), and then fit into the density with Isolde (Croll 2018).

AlphaFold2 (Jumper, Evans et al. 2021) was also used to generate structural predictions of the *L. interrogans* proteins FlaB1, FlaB2, FlaB3, and FlaB4. As with the *L. biflexa* samples, each FlaB isoform was individually fit into the density with Isolde (Croll 2018). The fit of several FlaB1-specific bulky side chains and the lack of density for specific bulky residues in the other isoforms suggested a majority FlaB1 population. As this flagellum (*flaA2^-^*) is symmetric, FlaB1 monomers were placed into each FlaB protofilament without additional Isolde refinements.

FcpA and FcpB have been crystallized in *L. biflexa* and *L. interrogans*, respectively (San Martin, Mechaly et al. 2017, Gibson, Trajtenberg et al. 2020). AlphaFold2-predicted structures of these proteins in *L. interrogans* closely resembled the crystal structures, and were used as an initial fit into the *flaA2*^-^ density. Isolde was then used to further refine the models.

The Isolde-modeled structures of the *L. biflexa* FlaB4 core, FlaA2, and FlaAP, as well as the *L. interrogans* FcpA and FcpB, were fit into the wild-type density using ChimeraX. Refinement of these models in the wild-type structure was not performed.

### Docking of the FlaA2 and FlaAP models into the wild-type density

The program Situs (Wriggers 2012, Kovacs, Galkin et al. 2018) was employed to dock our FlaA2 and FlaAP models into the wild-type density. The inner-curvature density (which remained unidentified in the tomographic wild-type structure) was used for the docking. A model consisting of one FlaA2 and one FlaAP monomer was used for the docking, which was performed with the program colores, using the inner core wild-type map, an angular sampling of 10°, and an anisotropy factor of 4. Manual docking of these sheath factors in the wild-type density and model, using a model consisting of one core repeat (11 protofilaments), one FlaA2 monomer, and one FlaAP monomer, was consistent with the Situs fittings.

## Acknowledgments

This work was supported by National Institutes of Health grants R01 GM110530 (to C. V. S.), T32GM8283 (to M.R.B) and U01 AI 088752, R01 TW009504, R01 AI052473, R0 AI121207 (to A. K.).; Agencia Nacional de Investigacion e Innovacion grant FCE_3_2016_1_126797 and Agence Nationale de la Recherche grant ANR-18-CE15-0027-01 (to A. B). Cryo-EM data was collected at the Yale CryoEM Resource that is funded in part by the NIH grant 1S10OD023603-01A1. We also thank the Center for Cellular and Molecular Imaging Electron Microscopy Facility at the Yale Medical School for assistance with this work and thank the Yale Center for Research Computing and the Yale High Performance Computing facility for guidance and use of the research computing infrastructure.

## Notes

### Competing Interest Statement

The authors have declared no competing interest.

