## Supplemental Figures and Tables for "Structural basis of flagellar filament asymmetry and supercoil templating by *Leptospira* spirochete sheath proteins"

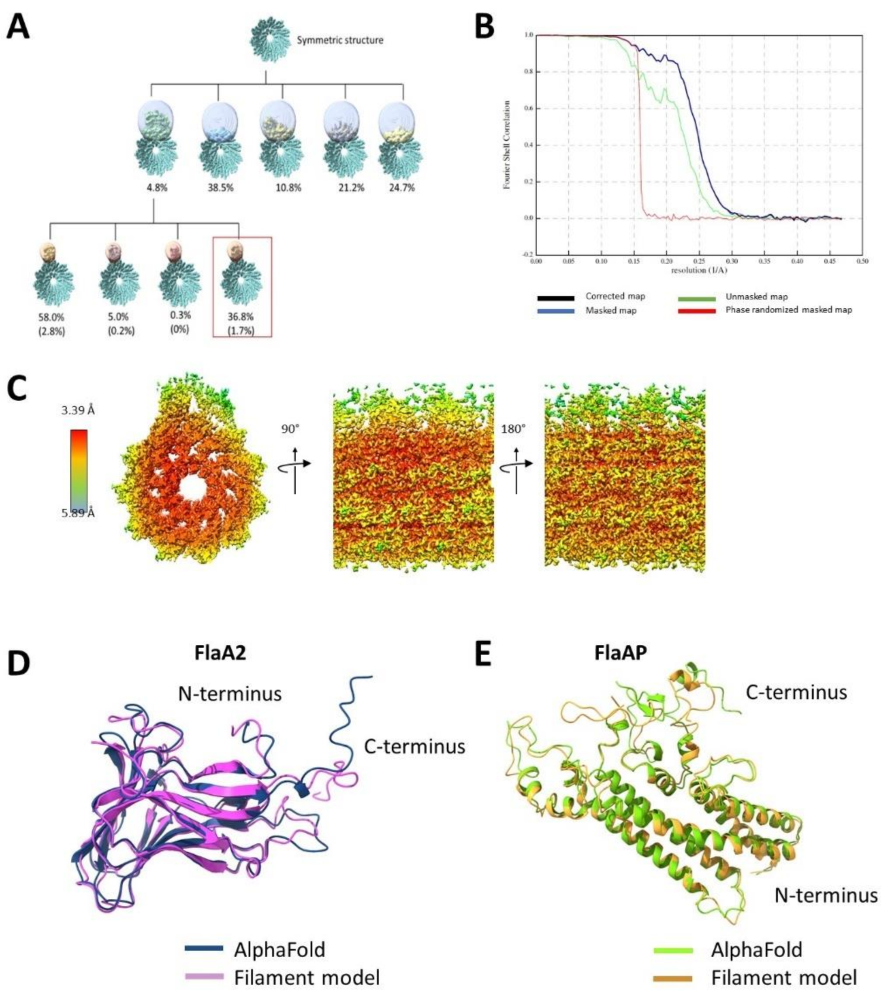


**Supplementary Figure 1.** FlaA2 and FlaAP are localized to the inner curvature of the asymmetric *fcpA*$^{-}$ flagellar filament. **A,** Focused classification of the *fcpA*$^{-}$ filaments. An initial refinement of the core was performed before masked classification was performed on the presumptive sheath region. The first round revealed three 'bare' classes and two sheathed classes (one with strong density within the mask, and one with density cut off by the mask). Another round of focused classification was performed on the sheathed class, using a mask focused on the beta-sheet sheath density. The best-resolved class (in the red rectangle) was used for all further analysis, and consisted of 1.7% of the initial particles. **B,** FSC curve of the *fcpA*$^{-}$ reconstruction, giving a resolution of 3.8 Å at an FSC of 0.143. The curve for the corrected map is shown in black, the curve for the masked map is shown in blue, the curve for the unmasked map is shown in green, and the curve for the phase randomized masked map is shown in red. **C,** Local resolution of the asymmetric filament, showing higher resolution was achieved in the core, and poorer resolution in the sheath regions. **D,** Comparison of the FlaA2 AlphaFold2 predicted model (in blue) and model fit into the density (in pink). Overall, the prediction is similar to the fit model at all locations except for the C-terminal region. **E,** Comparison of the FlaAP AlphaFold2 predicted model (in green) and fit model (in orange), showing very good agreement in the helices and most of the loop regions.

**A**

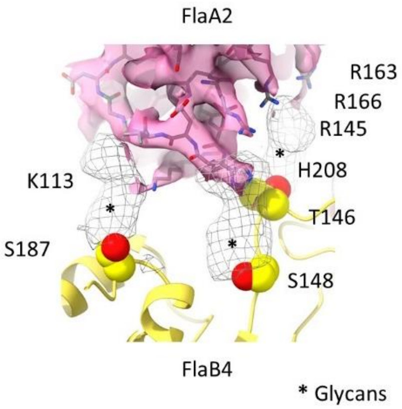


**Supplementary Figure 2.** Modeled FlaA2 and FlaAP sheath proteins fit well into the density **A,** The FlaA2 side chains fit well into the sheath density, with the placement of several positively charged residues closely interacting with the core glycans strongly suggested by cryo-EM density (shown in black mesh).

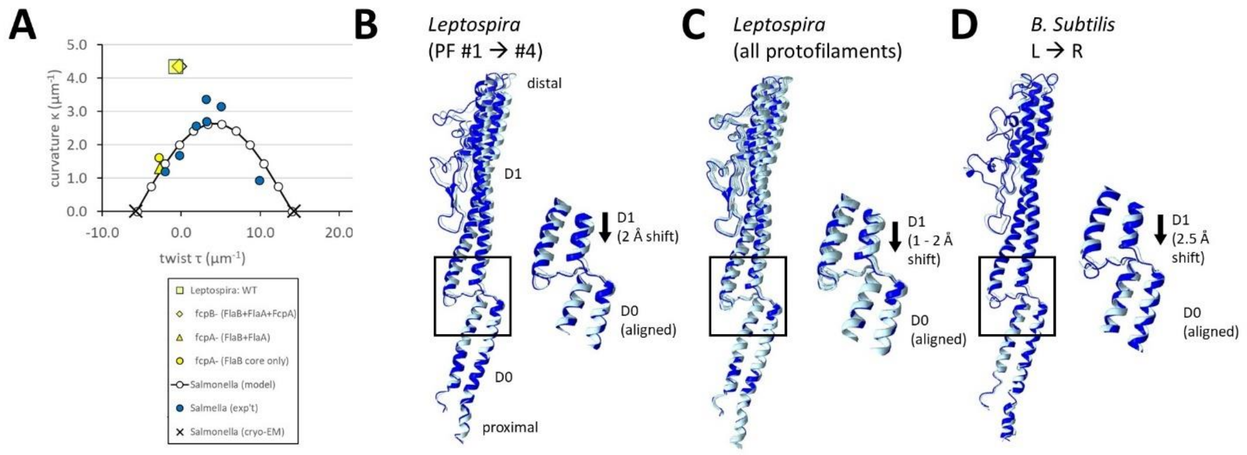


**Supplementary figure 3.** The core of the *fcpA*- structure resembles a 10L/1R flagellum. **A,** Comparison of the curvature and twist values of *Leptospira* and *Salmonella*. Values for the *Leptospira* wild-type, $fcpA^-$ (both sheathed and core-only), and $fcpB^-$ filaments were calculated, and are represented by a yellow square, triangle, circle, and diamond, respectively. Experimental values from *Salmonella* are shown as blue circles, values from cryo-EM models of *Salmonella* are shown as X's, and values from the Calladine model (Calladine 1975, Calladine, Luisi et al. 2013) are shown as white circles. **B,** Comparison of the models of protofilament #1 and protofilament #4. Insert on the right is a zoomed-in view of the D0-D1 linkers, highlighting the ~2 Å lateral shift of protofilament #4. Each monomer is aligned to the D0 domain. **C,** Comparison between all 11 individually modeled FlaB4 protofilaments, as in B. **D,** Comparison between the *B. subtilis* L and R protofilaments, revealing a ~2.5 Å lateral shift of 'R' relative to 'L', resembling the shift seen by the *Leptospira* protofilament #4.

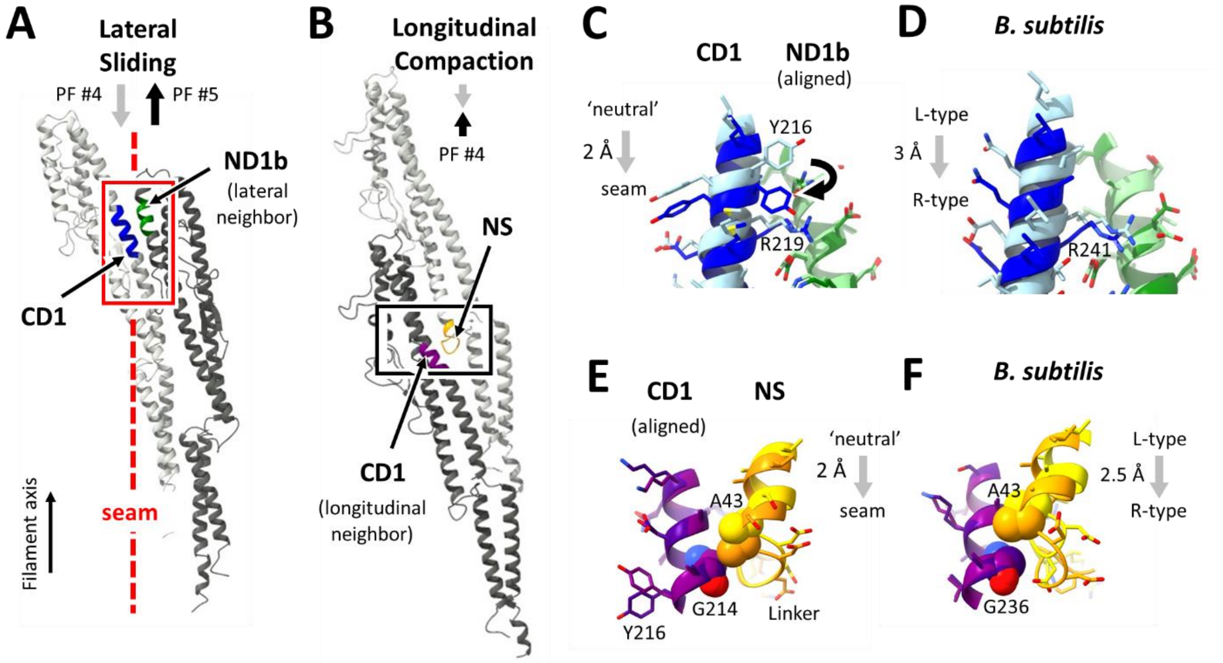


**Supplementary figure 4.** Structural signatures for the polymorphic switch mechanism found at the seam in the *fcpA⁻* structure. **A,** Overview showing a region of contact between FlaB D1 domains of two adjacent protofilaments, where lateral sliding occurs at the seam. The contact between CD1 and ND1b is shown in dark blue and dark green. **B,** Overview of two longitudinally adjacent FlaB subunits from a single protofilament. Within the protofilament, the CD1 helix interacts with the D0/D1 linker (NS) of the next proximal monomer in the row (boxed region). **C**. Comparison of seam (protofilaments #4-#5) and non-seam (protofilaments #11-#1) features, zoomed in to the boxed red region of **A**. Structures are aligned by superimposing their ND1b segments, revealing sliding of CD1 comparing the two protofilament pairs. Additionally, $Tyr_{216}$ of protofilament #4 undergoes a conformational change. **D**, Reference comparison of the transition between an 'all-L' filament (light blue and light green) to the 'all-R' filament from *B. subtilis* (dark blue and dark green; (Wang, Burrage et al. 2017)). Note the similar shift as compared to the one we observed in the *Leptospira* core, shown in panel **C**. **E,** The alignment between highlighted regions in **B**, protofilament #1 (light purple and yellow), and protofilament #4 (dark purple and orange). Structures are aligned to the CD1 helix. Protofilament #4 undergoes an axial shift, allowing $Ala_{43}$ of one monomer to contact $Gly_{214}$ of the next. **F,** As in E, but comparing this interaction in the 'all-L' protofilament (light purple and yellow) to the 'all-R' protofilament (dark purple and orange), showing analogous sliding of the NS domain.

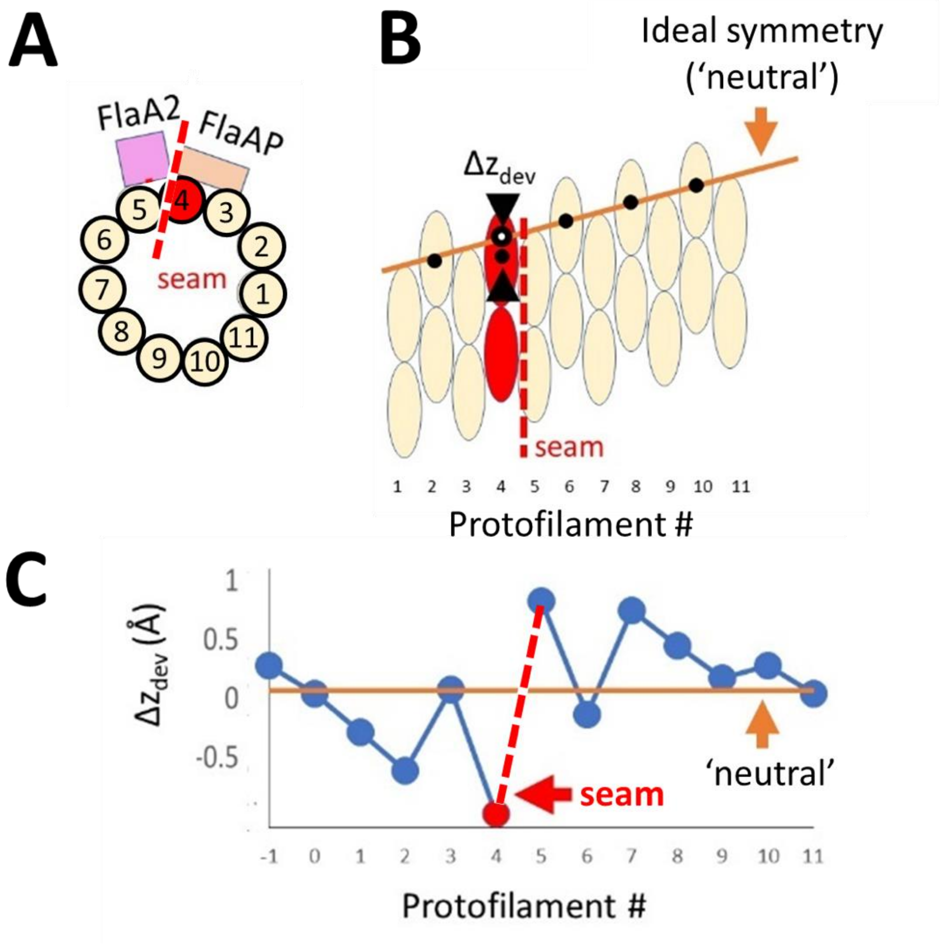


**Supplemental Figure 5.** Lateral sliding of protofilament #4 leads to a break in the helical symmetry at a 'seam'. **A,** Diagram showing the numbered protofilaments, as well as the placement of the FlaA2 and FlaAP sheath proteins. Protofilament #4 is colored red, and the seam is marked by a dashed line. **B,** Longitudinal view of two repeats of the FlaB core. Numbering and coloring as in A. The orange line represents what a helically symmetric filament would look like; deviations from this value ($\Delta z_{dev}$) are shown for protofilament #4 as the difference between the observed center of mass (black and white open circle) and the symmetric center of mass (black filled circle).**C,** A graph of $\Delta z_{dev}$ at each protofilament. This value is the difference between the observed center of mass z values, and the values that would be expected for a helically symmetric filament.

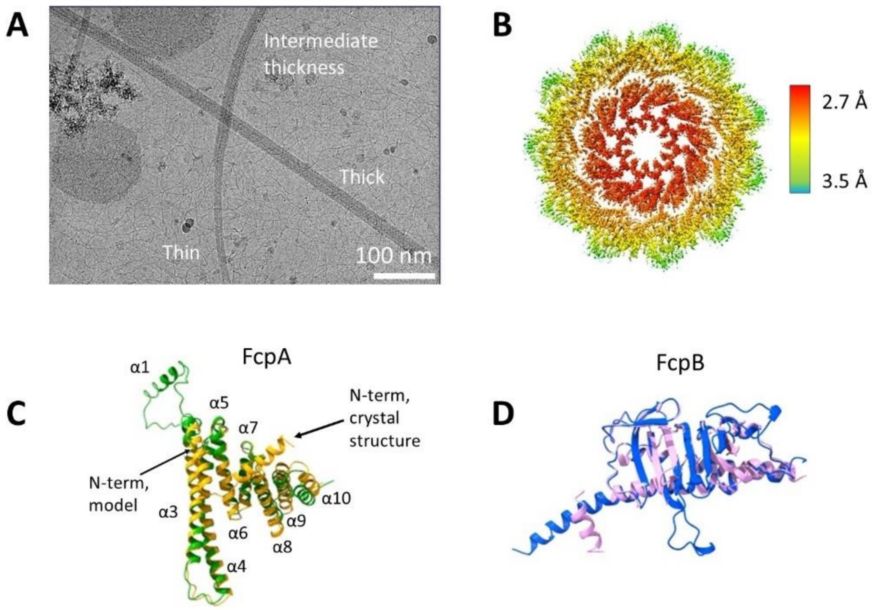


**Supplemental Figure 6.** A subset of the *flaA2*⁻ flagellar filaments are symmetrically decorated with FcpA and FcpB. **A,** Micrograph showing the heterogeneity in thickness and curvature of the *flaA2*⁻ filaments. The straight, thick filaments were analyzed structurally. **B,** Local resolution of the symmetric *flaA2*⁻ reconstruction. Highest resolution was achieved in the core, with slightly lower resolution in the sheath regions. **C,** Comparison of FcpA crystal structure (in orange) and the fit model (in green), highlighting a domain swap of the N-terminal helix α1. **D,** Comparison of the FcpB crystal structure (in pink) to the cryoEM structure (in blue).

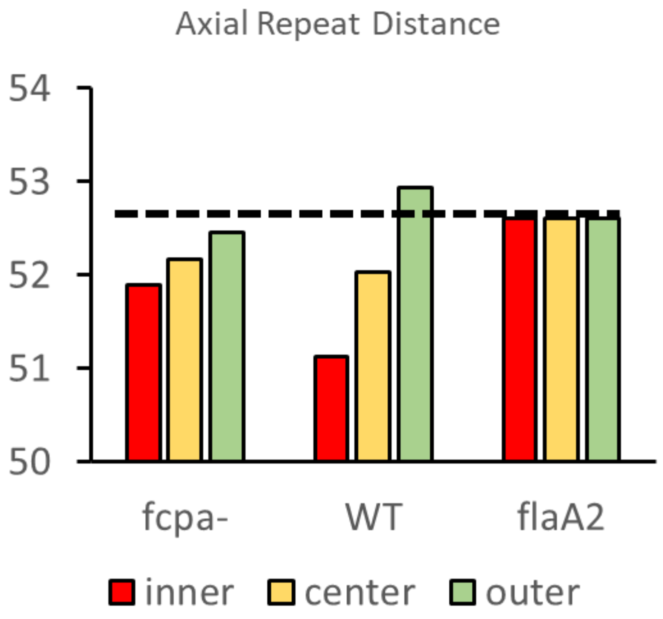


**Supplementary Figure 7.** FlaB axial repeat distances for wild-type and mutant *Leptospira* flagellar filaments, measured separately at inner and outer curvature locations. 'Center' value reflects the average of measured inner and outer values. Differences in the values for the three positions reflects the degree of curvature for each filament type, with wild-type being the most curved, followed by *fcpA*$^-$, while the *flaA2*$^-$ model is completely straight (helically symmetric). The FlaA2/FlaAP-sheathed *fcpA*$^-$ filaments have a 0.6 Å difference in the inner and outer curvature spacing. The tightly coiled wild-type filaments have a 1.8 Å difference in spacing between the inner and outer curvatures. The *flaA2*$^-$ filaments are symmetric and straight; the repeat spacing in these filaments is ~0.5 Å larger than the average spacing of the wild-type and *fcpA*$^-$ mutant filaments.

| | FlaB4 (1 repeat) | FlaA2 | FlaAP | FlaB1 (1 repeat) | FcpA | FcpB |
|---|---|---|---|---|---|---|
| **Clashscore, all atoms** | 1.08 ($99^{th}$ percentile) | 1.2 ($99^{th}$ percentile) | 2.27 ($99^{th}$ percentile) | 0 ($100^{th}$ percentile) | 0 ($100^{th}$ percentile) | 0.5 ($99^{th}$ percentile) |
| **Poor rotamers** | 0 | 0 | 0 | 0 | 0 | 0 |
| **Favored rotamers** | 99.18% | 98.9% | 98.94% | 99.58% | 100% | 99.55% |
| **Ramachandran outliers** | 0 | 0 | 0 | 0 | 0 | 0 |
| **Ramachandran favored** | 97.49% | 93.53% | 89.46% | 99.64% | 98.06% | 96.67% |
| **Rama distribution Z-score** | -2.23 ± 0.12 | -2.31 ± 0.49 | -4.43 ± 0.38 | 0.54 ± 0.44 | -0.70 ± 0.45 | 0.45 ± 0.53 |
| **MolProbity score** | 0.91 ($100^{th}$ percentile) | 1.26 ($99^{th}$ percentile) | 1.57 ($93^{rd}$ percentile) | 0.5 ($100^{th}$ percentile) | 0.5 ($100^{th}$ percentile) | 0.88 ($100^{th}$ percentile) |
| **CB deviations >0.25** | 11 (0.37%) | 2 (1.05%) | 8 (2.63%) | 0 | 0 | 0 |
| **Bad bonds** | 0 | 0 | 0 | 0 | 0 | 0 |
| **Bad angles** | 158 (0.48%) | 8 (0.34%) | 23 (0.63%) | 121 (0.40%) | 5 (0.17%) | 9 (0.33%) |
| **Cis prolines** | 0 | 0 | 0 | 0 | 0 | 0 |
| **CaBLAM outliers** | 15 (0.5%) | 8 (4.0%) | 19 (6.1%) | 22 (0.7%) | 2 (0.8%) | 2 (0.8%) |
| **CA geometry outliers** | 7 (0.23%) | 5 (2.51%) | 5 (1.62%) | 11 (0.36%) | 0 | 2 (0.84%) |

**Supplementary Table 1.** Molprobity (Chen, Arendall et al. 2010) was used to analyze the modeled FlaB4, FlaA2, FlaAP, FlaB1, FcpA, and FcpB structures.

| Locus | #Spectra Replicate 1 | #Spectra Replicate 2 | #Spectra Replicate 3 | ∑ #Spectra | Replicate Count | Description |
|---|---|---|---|---|---|---|
| tr\|B0SQZ5\|B0SQZ5_LEPBP | 675 | 190 | 246 | 1111 | 3 | FlaB4 |
| tr\|B0SSZ5\|B0SSZ5_LEPBP | 535 | 203 | 269 | 1007 | 3 | FlaB1 |
| tr\|B0SS97\|B0SS97_LEPBP | 177 | 241 | 225 | 643 | 3 | Iron(III) dicitrate TonB-dependent receptor putative signal peptide OS=Leptospira biflexa serovar Patoc fecA |
| tr\|B0SJC6\|B0SJC6_LEPBP | 322 | 138 | 163 | 623 | 3 | FlaAP |
| tr\|B0SSZ4\|B0SSZ4_LEPBP | 322 | 93 | 128 | 543 | 3 | FlaB2 |
| tr\|B0SLP3\|B0SLP3_LEPBP | 173 | 146 | 58 | 377 | 3 | Outer membrane protein OmpL1 putative signal peptide OS=Leptospira biflexa serovar Patoc ompL1 |
| tr\|B0SKT5\|B0SKT5_LEPBP | 182 | 92 | 99 | 373 | 3 | FlaA2 |
| tr\|B0SM66\|B0SM66_LEPBP | 145 | 83 | 104 | 332 | 3 | 30S ribosomal protein S2 OS=Leptospira biflexa serovar Patoc rpsB |
| tr\|B0SKT4\|B0SKT4_LEPBP | 210 | 67 | 46 | 323 | 3 | FlaA1 |
| tr\|B0SRP4\|B0SRP4_LEPBP | 129 | 106 | 85 | 320 | 3 | Putative cell envelope biogenesis protein putative signal peptide OS=Leptospira biflexa serovar Patoc LEPBI_I1733 |

**Supplementary Table 2.** The ten most abundant proteins identified from LC-MS/MS mass spectrometric analyses of purified *fcpA*$^{-}$ flagella. Besides FlaAP, additional uncharacterized proteins were identified with lower abundance (not shown).
